# Integration of system phenotypes in microbiome networks to identify candidate synthetic communities: a study of the grafted tomato rhizobiome

**DOI:** 10.1101/2019.12.12.874966

**Authors:** Ravin Poudel, Ari Jumpponen, Megan M. Kennelly, Cary Rivard, Lorena Gomez-Montano, Karen A. Garrett

## Abstract

Understanding factors influencing microbial interactions, and designing methods to identify key taxa, are complex challenges for achieving microbiome-based agriculture. Here we study how grafting and the choice of rootstock influence root-associated fungal communities in a grafted tomato system. We studied three tomato rootstocks (BHN589, RST-04-106 and Maxifort) grafted to a BHN589 scion and profiled the fungal communities in the endosphere and rhizosphere by sequencing the Internal Transcribed Spacer (ITS2). The data provided evidence for a rootstock effect (explaining ~2% of the total captured variation, p < 0.01) on the fungal community. Moreover, the most productive rootstock, Maxifort, supported greater fungal species richness than the other rootstocks or controls. We then constructed a phenotype-OTU network analysis (PhONA) using an integrated machine learning and network analysis approach based on sequence-based fungal Operational Taxonomic Units (OTUs) and associated tomato yield data. PhONA provides a graphical framework to select a testable and manageable number of OTUs to support microbiome-enhanced agriculture. We identified differentially abundant OTUs specific to each rootstock in both endosphere and rhizosphere compartments. Subsequent analyses using PhONA identified OTUs that were directly associated with tomato fruit yield, and others that were indirectly linked to yield through their links to these OTUs. Fungal OTUs that are directly or indirectly linked with tomato yield may represent candidates for synthetic communities to be explored in agricultural systems.

**IMPORTANCE:** The realized benefits of microbiome analyses for plant health and disease management are often limited by the lack of methods to select manageable and testable synthetic microbiomes. We evaluated the composition and diversity of root-associated fungal communities from grafted tomatoes. We then constructed a phenotype-OTU network analysis (PhONA) using these linear and network models. By incorporating yield data in the network, PhONA identified OTUs that were directly predictive of tomato yield, and others that were indirectly linked to yield through their links to these OTUs. Follow-up functional studies of taxa associated with effective rootstocks, identified using approaches like PhONA, could support the design of synthetic fungal communities for microbiome-based crop production and disease management. The PhONA framework is flexible for incorporation of other phenotypic data and the underlying models can readily be generalized to accommodate other microbiome or other ‘omics data.

## Introduction

Interactions are key to defining system behaviors, structures, and outcomes. In microbial systems, interactions among organisms define their distribution, assemblies, and ecosystem functions. In addition to microbe-microbe interactions, microbes interact with their hosts, and are essential to host health and performance (1–8). In agriculture, plant–microbe interactions improve plant productivity by providing access to nutrients (9–11), reducing infection by plant pathogens (5, 12), triggering plant growth promoting factors (13, 14), and enhancing plant resistance (15, 16) and tolerance to abiotic stresses (17–19). Although the importance of microbes and host-microbe interactions to host health and ecological processes is well-known, interaction-based approaches to manage crop-production remain a scientific frontier. Past attempts to translate information about microbial interactions to design biocontrol agents or biofertilizers have often had limited efficacy and durability (20, 21). Most microbial inocula have been applied as single species, often selected based on pairwise relations of microbes with a pathogen or the host. Interactions among microbes as well as with the host are important, and the net outcome of these complex interactions defines host health and ecosystem functions (22). Thus, it is critical to understand the ecology of microbes selected for biological applications, and systems approaches centered on host-microbe interaction can help guide the selection of microbes for synthetic communities (23).

Among tools to better understand microbial interactions, network models of microbial communities, and studies of network structures and key groups, have proven popular for generating hypotheses about how to engineer microbial consortia. In such network models of microbiomes, a node represents an OTU, and a link exists between two OTUs if their sequence proportions are significantly associated across samples. When evaluated with other conventional measures of microbial community structures, such as diversity indices, network models can be used to identify hub taxa that may be key to maintaining microbial assemblages and diversity (24), or to evaluate changes in community complexity and interactions in response to experimental treatments (25). Microbiome networks are useful for describing general community structures and their key properties and are often the most practical option when additional information about species interactions is missing or the goal is to compare across studies with different types of data (26, 27). The utility of network analysis for identifying candidate assemblages for biocontrol can be enhanced by incorporating nodes that represent other additional types of features (28, 29). For instance, a novel association of host metadata with the microbiome was revealed in an integrated microbiome-metadata network (30), where a feature strongly associated with hub microbes can serve as a marker to measure host performance. In agriculture, plant yield or other phenotypic traits can be integrated in microbiome networks, with the potential to identify microbial consortia that are predictive of host phenotypes. Because such models include host phenotypes, they facilitate finding candidate sets of OTUs that may directly or indirectly affect host phenotypic traits. Visualization of networks is often valuable for this purpose, but the real value of phenotype-based network models is their ability to infer potential candidate taxa or consortia. The hypothesized beneficial sets of OTUs may represent targets for pure culturing efforts or, if cultures exist, the sets can be further evaluated in laboratory or field studies.

While phenotype-based network models have the potential to identify key taxa, application of such models should be integrated with findings from other community analyses so that the inference about key taxa is biologically and ecologically meaningful. For example, plant microbiome studies indicate that a small but consistent proportion of variation in microbial communities is often explained by the host genotype (31–37), indicating the potential for genotype-based modulation of microbial communities in crop-production on a broader scale. These results support the idea of host-specific microbial community selection (38). Many such microbes may be taxa that are evolutionarily essential for the survival and function of plants (39, 40). In addition, the extent of host genotype filtering of microbes differs across the rhizosphere, rhizoplane, and endosphere, and varies from one host species to another (41–43). Results that indicate microbial filtering by different crop hosts, plant compartments, geographic locations, and environmental factors (44, 45) are promising for designing experiments to minimize the search space, or necessary sample numbers, to identify candidate taxa for synthetic communities. For instance, factors that explain great variation in microbial community composition, but that are outside the control of management, can be treated as blocks in experimental designs, so that host- or compartment-specific effects on the microbial community can be searched to identify the most desirable candidate taxa.

In our current study, building on previously described agricultural experiments in grafted tomato systems (46, 47), we characterized the root-associated fungal (RAF) communities and implemented an interaction-based approach to select potential candidate fungi that are predictive of tomato yield and/or that are in significant association with other fungal taxa. The new phenotype-OTU network analysis (PhONA) is a method for network-visualization and a framework to support the selection of candidate taxa and to integrate system traits (such as host yield) in microbe-microbe association networks. PhONA first identifies OTUs predictive of phenotype using lasso regression, then uses the predicted OTUs from lasso regression to build a reduced GLM. PhONA then combines the GLM results indicating positive or negative associations of the predicted OTUs with the host phenotype as well as with other OTUs in a network model (Fig. 1). Due to the large number of OTUs compared to the sample size, lasso regression was used because it is suited for minimizing overfitting when applied with a relatively small sample size (48) and has been implemented in microbiome studies (49, 50).

**FIG 1.**
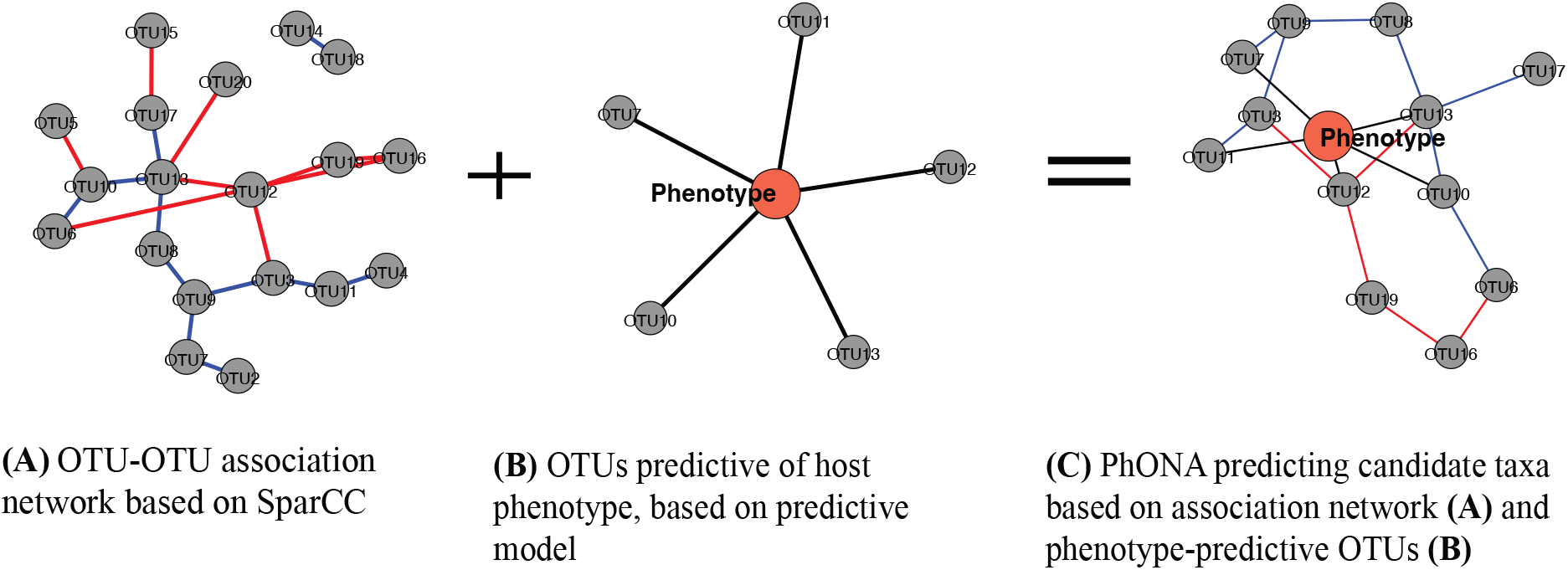
The Phenotype-OTU network analysis (PhONA) combines (A) an OTU-OTU association network with (B) the nodes selected based on predictive model for their association with a host phenotype variable such as yield, to create (C) a PhONA.

Phenotype-based selection of microbial consortia is promising as an effective approach to select representative microbial taxa and could support the design of microbiome-based products. Changes in abundance (51), successive selection over multiple generations (3), or analyses of binary host-microbe relationships (52) are some of the recent phenotype-based applications to select candidate taxa for biological applications. Despite the importance of biological test-based approaches, difficulty in culturing all the microbes makes computational approaches instrumental to define microbe-microbe and host-microbiome associations, and to identify the biological and ecological key taxa. Tools to describe the community structures based on the co-occurrence matrix or covariance structures (53, 54) are more common, whereas tools to integrate host phenotype or environmental factors are at an earlier phase of development. Relatively small sample sizes combined with large number of features may limit applications of the recent graph-based approaches. Such methods allow measurement of direct associations via conditional dependence structures and offer options to include environmental and phenotypic information in the model (55). CoNet (56) and Flashweave (55) allow representation of the phenotype or an environmental variable as an extra node or a column in the adjacency matrix, and the same statistical method can be applied to define the associations among microbes and between microbes and phenotypes (taxa and metadata). PhONA is generic as it allows the user to select data structure-specific models for microbe-microbe and microbe-phenotype associations.

In the current case study, we used lasso regression to identify the subset of OTUs and then fitted them using GLMs to predict OTU-phenotype associations, whereas the OTU-OTU associations were defined using SparCC. Additionally, we contrasted the RAF community’s diversity and interactions among the rootstocks and the controls, for endosphere and rhizosphere compartments. Based on our yield data, rootstock vigor, and previous studies of microbial interactions (25), we expected a greater number of fungal OTUs and of microbial associations for more productive rootstocks. Moreover, in our previous studies of bacterial communities in the tomato rhizobiome, we observed compartment-specific (endosphere vs rhizosphere) effects of grafting and rootstocks on bacterial community composition and diversity (47), and expected similar effects on RAF diversity and composition. All the code and vignettes for PhONA are available at https://ravinpoudel.github.io/PhONA/index.html and archived at zenodo (DOI: 10.5281/zenodo.6600986).

## METHODS

### Experimental Plots, Rootstocks, and Study Sites

We studied grafted tomato plants in high tunnels in an experimental design similar to that described by Poudel et al. (47). Tomato plants were grafted following a tube-grafting protocol described in Meyer et al. (46). Our study included three rootstocks (BHN589; RST-04-106, and Maxifort) in four graft treatments: 1) nongrafted BHN589 plants; 2) selfgrafted BHN589 plants (plants grafted to their own rootstock); 3) BHN589 grafted to RST-04-106; and 4) BHN589 grafted to Maxifort. We chose BHN589 as scion primarily based on its popularity due to high yield and high-quality fruit with a long shelf life. For rootstocks, we selected Maxifort because it is a productive and popular rootstock, and RST-04-106 as a new rootstock variety based on tomato breeders’ recommendations.

Our study included two sites: Olathe Horticulture Research and Extension Center (OHREC) and Common Harvest Farm, a farm managed by a collaborating farmer. For more information about the sites, see Table S1. At each study site, the four graft treatments were assigned to four plots per block in a randomized complete block design. Each plot consisted of 5-8 plants, and one middle plant per plot was sampled during the peak growth stage. There were six blocks at OHREC, and four blocks at Common Harvest Farm, such that for each year, each graft treatment was replicated 10 times. The experiment was repeated for two years (2014 and 2015) with a similar design, with the blocks and rootstocks randomly and independently assigned in each year.

### Sample Preparation, DNA Extraction, and Amplicon Generation

To compare the fungal communities, we selected a center plant from each plot and carefully dug the whole plant out such that the majority of the roots remained intact. Endosphere and rhizosphere samples were prepared as previously described (47) and the total genomic DNA was extracted using a DNA extraction kit (MoBio UltraClean Soil DNA Isolation Kit; MoBio, Carlsbad, CA, USA) as per manufacturer’s protocol, with slight modification during the homogenization step (47). To recover high genetic diversity, we opted for the two-step PCR approach. The primary PCR amplicons were generated in 50 μL reactions under the following conditions: 1 μM forward and reverse primers, 10 ng template DNA, 200 μM of each dioxynucleotide, 1.5 mM MgCl2, 10 μL 5x Phusion Green HF buffer (Finnzymes, Vantaa, Finland), 24.5 μL molecular biology grade water, and 1-unit (0.5 μL) Phusion Hot Start II DNA Polymerase (Finnzymes, Vantaa, Finland). PCR cycle parameters consisted of a 98° C initial denaturing step for 30 seconds, followed by 30 cycles at 98° C for 10 seconds, 57° C annealing temperature for 30 seconds, and 72° C extension step for 30 seconds, followed by a final extension step at 72° C for 10 minutes. All samples were PCR-amplified in triplicate to minimize stochasticity, pooled, and cleaned using Diffinity RapidTip (Diffinity Genomics, West Chester, PA, USA). In this PCR, we amplified the entire ITS region of fungal rRNA genes using primers ITS1F-CTTGGTCATTTAGAGGAAGTAA and ITS4-TCCTCCGCTTATTGATATGC (<u>e.g.</u> 57). The average amplicon length of the ITS region in fungi is about 600 bp and could not reliably be fully covered with the Illumina MiSeq platform (v.3-chemistry) in a single read. Thus, in the following nested PCR, only ITS2 of the ITS region was amplified using fITS7-ITS4 primers (58) incorporating unique Molecular Identifier Tags (MIDs) at the 5’ end of the reverse primer (ITS4). For the nested PCR, we used similar reagents and PCR conditions as in the primary PCR, with some modifications: the number of PCR cycles was reduced to ten, total reaction volume was reduced to 25 ul, and 5 ul of cleaned PCR product from the first PCR amplification was used as the DNA template. The nested PCR was also run in triplicate, pooled by experimental unit, and cleaned with an Agencourt AmPure cleanup kit using a SPRIplate 96-ring magnet (Beckman Coulter, Beverly, MA, USA) as per the manufacturer’s protocol. Then, 200 ng of cleaned, barcoded amplicons were combined per experimental unit, and the final pool was cleaned again using an Agencourt AmPure cleanup kit as above. Illumina MiSeq adaptors were ligated to the library and paired-end sequenced on a MiSeq Personal Sequencing System (Illumina, San Diego, CA, USA) using MiSeq Reagent Kit V3 with 600 cycles. The endosphere and the rhizosphere amplicon libraries were sequenced separately in two runs. Adaptor ligation and sequencing were performed at the Integrated Genomics Facility at Kansas State University. All sequence data generated in this study were deposited in the NCBI Sequence Read Archive depository (BioProject:…….).

### Bioinformatics and OTU Designation

The sequence library of fastq files was curated using the MOTHUR pipeline (Version 1.33.3; (59)) following steps modified from the MiSeq Standard Operating Protocol (SOP; www.mothur.org/wiki/MiSeq_SOP). Briefly, the forward and the reverse reads were assembled into contigs using the default alignment algorithm. Any sequences shorter than 250 base pairs or containing an ambiguous base call or more than eight homopolymers or missing MIDs were removed from the library. Barcoded sequences were assigned to experimental units, and the data for endosphere and rhizosphere libraries were merged and processed together for the remaining steps in the MOTHUR pipeline. The pairwise distance matrix based on the filtered sequences was created and sequence data clustered into OTUs at 97% sequence similarity using the nearest neighbor joining algorithm. The clustered OTUs were assigned to a putative taxonomic identity using a Bayesian classifier (60) referencing the UNITE plus INSD non-redundant ITS database (61). To minimize the bias resulting from unequal sequence counts per sample, samples were rarified to the lowest sequence count among the samples (6,777). The final curated OTU database included 1,084,281 total sequences representing 16,151 fungal OTUs, including singletons (5,376).

### Statistical analyses

We evaluated the network of associations among fungal OTUs with network models to better understand the community composition and the interactions therein. The observed OTU database was divided into eight subsets, each combination of the four rootstocks and two compartments (endosphere and rhizosphere), such that we constructed eight networks in total. In our network models, a node represents an OTU and a link exists between a pair of OTUs if there is evidence (p < 0.05) that their frequencies are correlated (positively or negatively) across samples. Reducing false associations due to compositional bias in network modeling of microbiome data is important for clearer interpretation (62). Thus, we used a Sparse Correlations for Compositional data (SparCC) method to evaluate the pairwise associations (62), designed to minimize the compositional bias effect due to normalization. In our analyses, associations were defined in 20 iterations, and the significance of a pairwise association was determined from 100 bootstrapped datasets. Once the matrix defining all the pairwise associations was derived, we selected only those OTUs for which the absolute value of at least one association was greater than 0.5 (and p < 0.05) in the network analyses for each of the rootstock genotypes.

To identify the OTUs associated with tomato yield in each rootstock, a regression-based model was fitted to the observed data. Marketable tomato yield data reported by Meyer et al. (46) was the response variable and fungal OTUs were potential predictors. We used the caret package (63) to evaluate the lasso regression and selected OTUs using varImp functions. Lasso regression used an L1 regularization approach to shrink the less important variables’ coefficients to zero and to reduce the number of variables in the model. In lasso regression, lambda determines the penalty of regularization, and its value can range from zero to infinity; when it is zero, the results are similar to the least square lines. A grid-based approach was used to tune the lambda parameter using repeated (ITERS=500) 5-fold cross-validation and the value of lambda with lowest variance was selected. Only the OTUs with non-zero coefficients were selected, based on the lasso-regression model, to build the reduced GLM model, and the association type of each OTU with phenotype was estimated. Given the small sample size, we did not evaluate the model performance by splitting the data into training and test cases, although this would be a valuable step in future studies with larger sample sizes. PhONA then integrates the results from the GLM model for yield with the OTU-OTU association network. We plotted the resulting network using the igraph package (54) in R. To evaluate the role of nodes in the network, we placed each node in one of four categories – peripherals, module hubs, network hubs, and connectors – based on the within-module degree and among-module connectivity (24, 55). Role analyses were only used the presence or absence of links in the network and do not account for the link types (i.e. positive or negative associations).

To evaluate the effects of rootstocks on fungal diversity, Shannon entropy and species richness were evaluated using the vegan package (64) wrapped by the phyloseq package (65) in R (66). Differences in diversity across the rootstocks were compared using a mixed model ANOVA in the lme4 package in R (67). Study site and sampling year were treated as random factors, blocked by study sites, whereas the rootstock and compartments were treated as fixed factors. Changes in fungal community composition across the samples were estimated based on a Bray-Curtis dissimilarity distance matrix and visualized in non-metric multidimensional scaling (NMDS) plots. The contribution of factors to the observed variation in fungal composition was estimated in a permutational multivariate analysis of variance (PERMANOVA, using 1000 permutations) using the adonis function in the vegan package (64). To identify OTUs that were sensitive to the rootstock treatments, the observed frequency (proportion) of each OTU was evaluated by fitting a generalized linear model (GLM) with negative binomial distribution, to identify depleted or enriched OTUs (Differentially Abundant OTUs – DAOTUs). Likelihood ratio tests and contrast analyses (between the hybrid rootstock and controls) were performed for the fitted GLM to identify the DAOTUs. We used OTU frequencies from selfgrafts and nongrafts as controls, in comparisons with other rootstocks, using contrasts. All tests were adjusted to control the false discovery rate (FDR, p < 0.01) using the Benjamini-Hochberg method (68). A differential abundance test was performed within the controls (selfgraft vs. nongraft) to identify the OTUs responsive to the grafting procedure, itself.

## RESULTS

### RAF in the Grafted Tomato System

Once rare OTUs (<10 sequence counts, which accounted for more than 90% of the observed OTUs) were removed, the community consisted of 1586 OTUs and 1,063,017 sequences. Of these sequences, 4.8% remained unclassified at the phylum level (Fig. S1). The classified sequences represented Ascomycota (52.5%), Basidiomycota (25.6%), Zygomycota (11.5%), Chytridiomycota (3.6%), Glomeromycota (1.7%), and Rozellomycota (0.07%) (Fig. 1). At the class level, Pezizomycetes, Agaricomycetes, and Dothideomycetes were the most abundant across all the rootstocks. At the order level, the communities were dominated by Pezizales, Pleosporales, Cantharellales, Mortierellales, and Hypocreales. Analyses at the family level revealed that Pyronemataceae, Mortierellaceae, Ceratobasidiaceae, and Pleosporaceae were the most common taxa overall. At the genus level, *Mortierella, Thanatephorus*, and *Alternaria* were the most abundant genera.

### Effects of Grafting and Rootstock on α-Diversity

There was strong evidence for a rootstock effect on OTU richness (F_*1, 3*_ = 8.6, p < 0.001) and Shannon entropy (F_*1, 3*_ = 3.2, p = 0.02) for tomato RAF communities. Mean species richness was higher in both the endosphere (p = 0.01) and rhizosphere (p = 0.001) of one of the hybrid rootstocks, Maxifort, compared to the nongrafted control. Shannon entropy followed trends similar to richness with a higher estimate for Maxifort; however, there was only evidence for higher Shannon entropy in Maxifort for the rhizosphere (p = 0.004), but not for the endosphere (p = 0.6) (Fig. 2). Both species richness and Shannon entropy were higher (p < 0.001) in the rhizosphere than in the endosphere across all the rootstock genotypes (Fig. S2).

**FIG 2.**
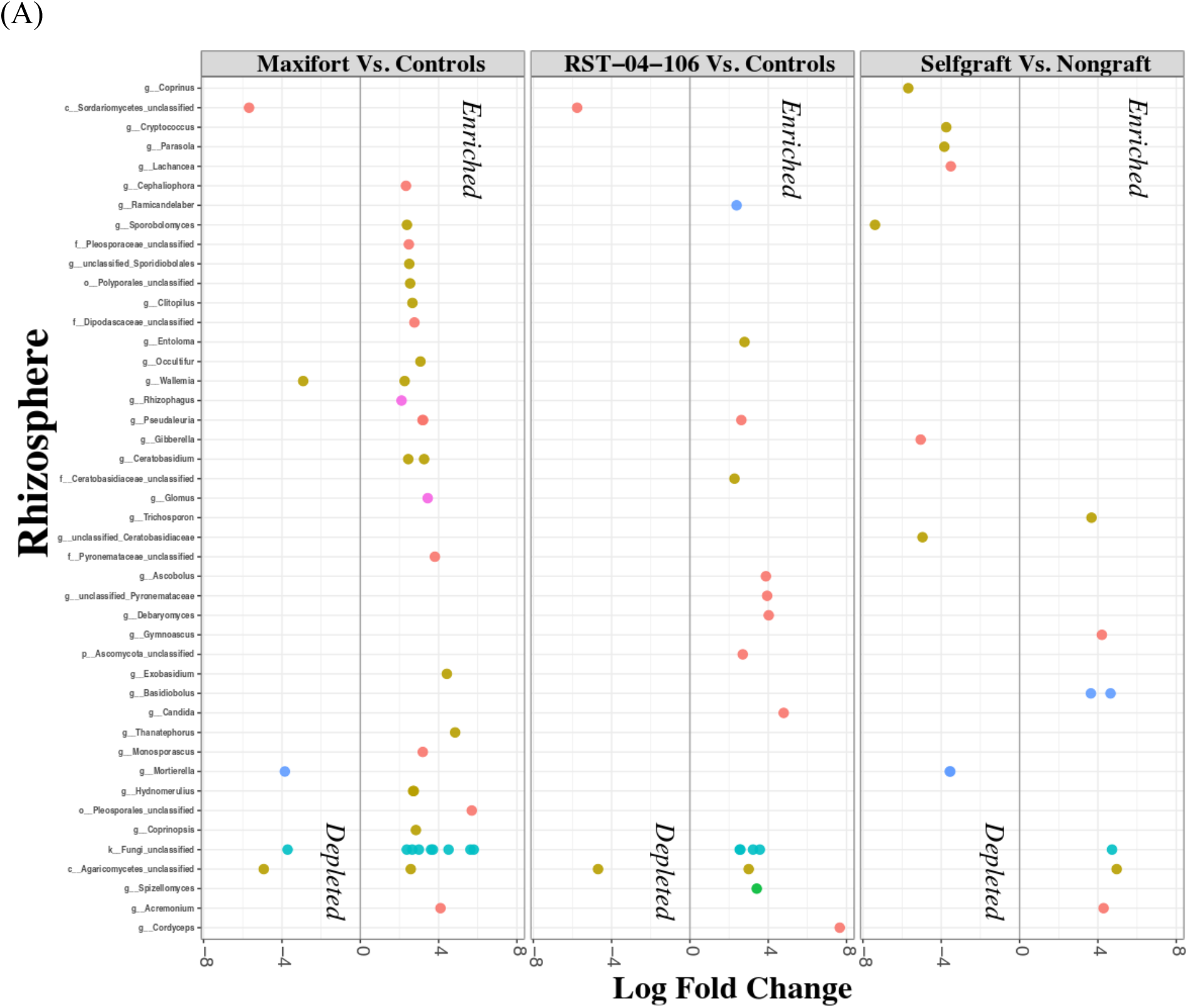
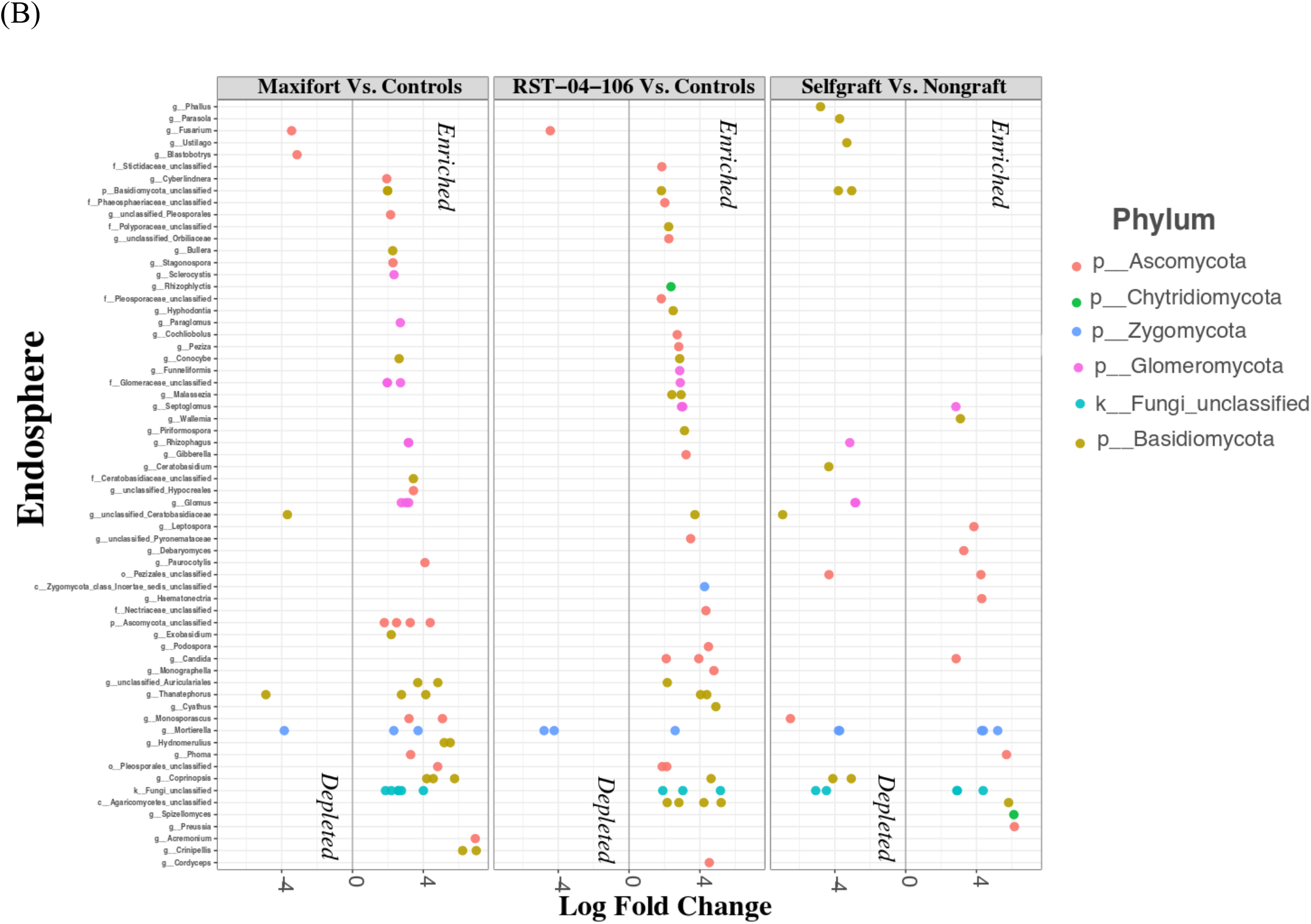
Enriched and depleted OTUs across tomato rootstock genotype combinations (nongraft BHN589, selfgraft BHN589, and BHN589 grafted on two hybrid rootstocks (RST-04-106 and Maxifort)) evaluated for the rhizosphere (A) and the endosphere (B), using OTU counts from selfgrafts and nongrafts as controls. All the tests were adjusted to control the false discovery rate (FDR, p < 0.01) using the Benjamini-Hochberg method. Each point represents an OTU labeled at the genus level and colored based on phylum, and the position along the x-axis represents the abundance fold-change contrast with controls (except for the selfgraft vs. nongraft comparison, where the nongraft treatment was used as a control for the contrast).

### Effects of Grafting and Rootstock on RAF Composition

Based on previous studies of the plant genotype effect on the rhizobiome (47), we expected a significant rootstock effect on community composition. Rootstock explained 2% of the variation in the RAF community composition (PERMANOVA; p<0.01), whereas compartment, study site, and year explained a greater proportion of the variation than rootstock (Fig. S3 and Table S2). Endosphere-rhizosphere compartments accounted for 8.92% of the variation. Study site and interannual variation explained 8.34% and 5.38% of the total variation, respectively (Table S2).

### Comparison of DAOTUs

The analysis of differential abundance found effects of rootstock genotype at the individual OTU level. While analyses of alpha diversity indicated higher diversity in the rhizosphere than in the endosphere, we observed the opposite in the analysis of DAOTUs, with nearly twice as many DAOTUs in the endosphere (n = 146 i.e. 9.2% of the total OTUs) compared to the rhizosphere (n = 76 i.e. 4.8% of total OTUs) (Figs. 2 and S4). Comparison across rootstocks indicated a greater number of DAOTUs in Maxifort (n = 80) than in RST-04-106 (n = 66) and the selfgraft control (n = 49). Compared to the hybrid rootstocks, the number of depleted taxa was greater in the selfgraft control (n = 28). Among the enriched OTUs in Maxifort, 27 OTUs belonged to Basidiomycota, 20 to Ascomycota, and 11 to Glomeromycota, whereas four basidiomycete, three ascomycete, and one zygomycete OTUs were depleted in Maxifort. In RST-04-106, enriched taxa included 22 OTUs in Basidiomycota, 20 OTUs in Ascomycota and five OTUs in Glomeromycota, whereas the depleted OTUs included six in Zygomycota. Comparing the self- and nongraft controls, nine OTUs in Ascomycota, three OTUs in Basidiomycota, and four OTUs in Zygomycota were enriched in the selfgraft treatment, whereas 12 OTUs in Basidiomycota, seven OTUs in Ascomycota, four OTUs in Zygomycota, and three OTUs in Glomeromycota were depleted in the selfgraft treatment.

### Network Analysis/ General Network Structures

Fungal community complexity, defined in terms of mean node degree and community structures/motifs, varied among the rootstocks in both the endosphere and the rhizosphere, with a greater mean node degree in one of the hybrid rootstocks, Maxifort, compared to both controls and RST-04-106 (Figs. 3, S5, S6, and Table S3). Complexity was higher in the rhizosphere than in the endosphere compartment (Figs. 3, S5, S6, and Table S3). In addition to the total number of links, the link type (either positive or negative) differed among the rootstocks in both compartments (Table S3), with a higher ratio of negative to positive links in Maxifort in both the endosphere and the rhizosphere compartments. Rhizosphere fungal communities had a higher ratio of negative to positive links than those in the endosphere, for all rootstocks. Although we observed rootstock-specific or compartment-specific effects on the node degree and ratio of negative to positive links, the number of modules defined using a simulated annealing (SA) algorithm were similar in both the endosphere and rhizosphere compartments and across the rootstocks (Table S3). Our analyses of node types divided the observed nodes in the association-network into four categories: peripherals, module hubs, network hubs, and connectors. More taxa in the rhizosphere were identified as key nodes than in the endosphere, and key nodes were more common in the hybrid rootstocks than in the non- and selfgrafted controls (Figs. 4 and 5).

**FIG 3.**
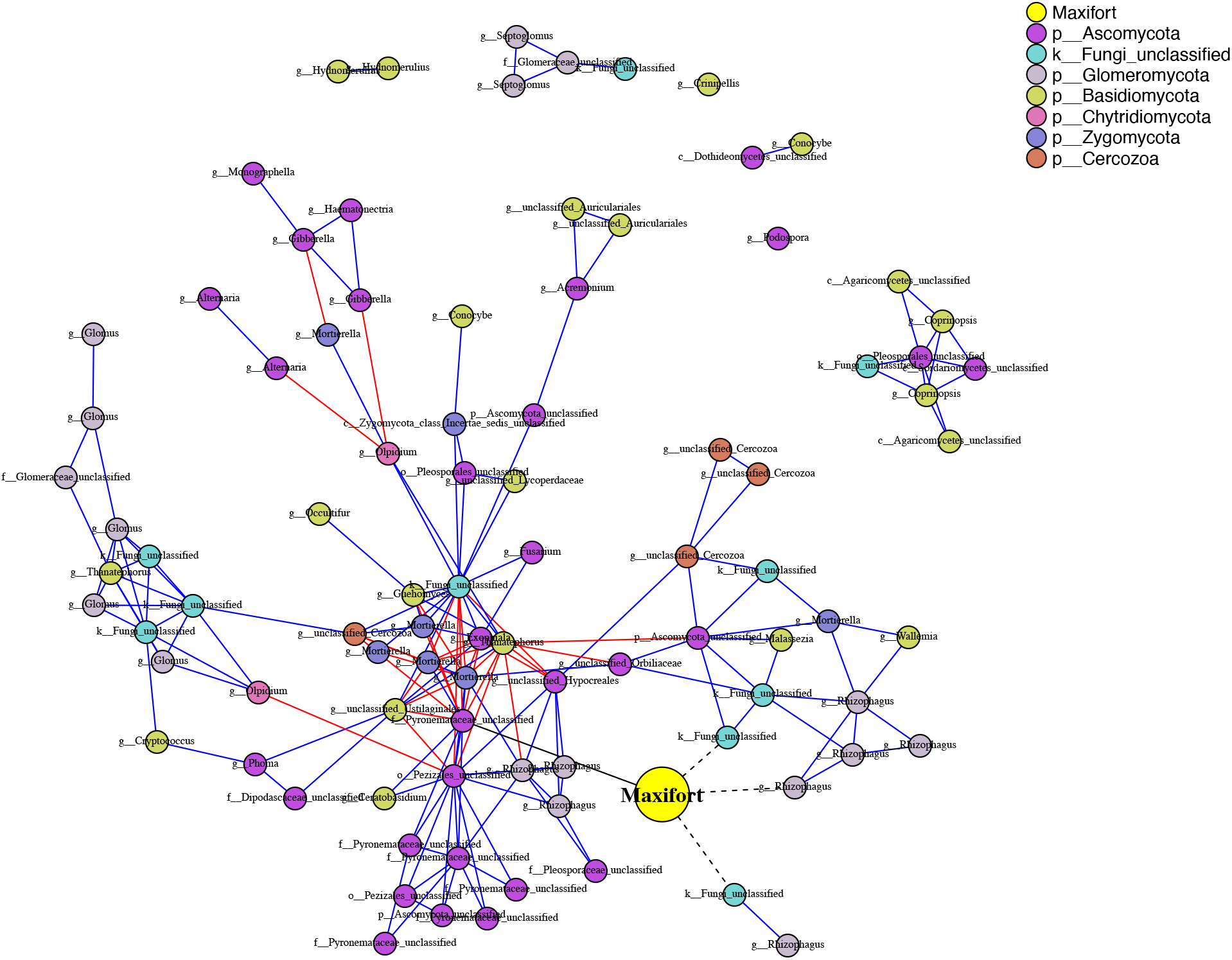
Phenotype-OTU network analysis (PhONA) of endosphere fungal taxa for BHN589 grafted on Maxifort. Node color indicates the phylum, except that the yellow-color node represents yield associated with the rootstock. Nodes connected to the rootstock yield node with black links are taxa that were predictive of rootstock yield, where dotted and solid lines indicate negative and positive associations with the yield node, respectively. Red and blue links represent negative and positive associations, respectively, between OTUs. Nodes are labeled with the finest-resolution taxonomic categorization available.

**FIG 4.**
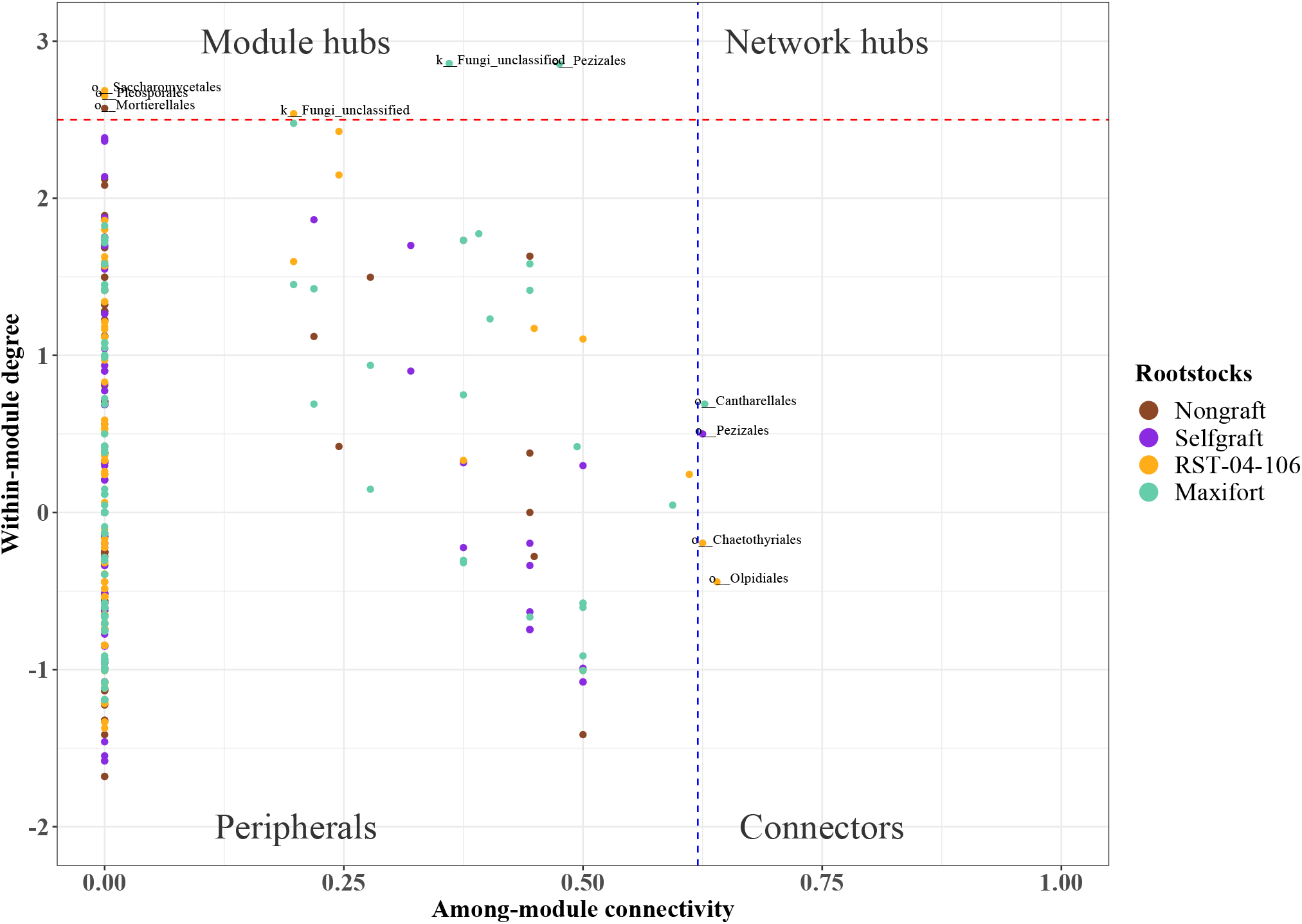
Partitioning of endosphere fungal OTUs according to their network roles. Nodes were divided into four categories based on within-module degree and among-module connectivity. The blue dashed line represents a threshold value (0.62) for among-module connectivity, and the red dashed line represents a threshold value (2.5) for within-module degree. Nodes were categorized as peripherals, connectors, module hubs, and network hubs. Node color indicates rootstock treatment (nongraft BHN589, selfgraft BHN589, and BHN589 grafted on two hybrid rootstocks (RST-04-106 and Maxifort)).

**FIG 5.**
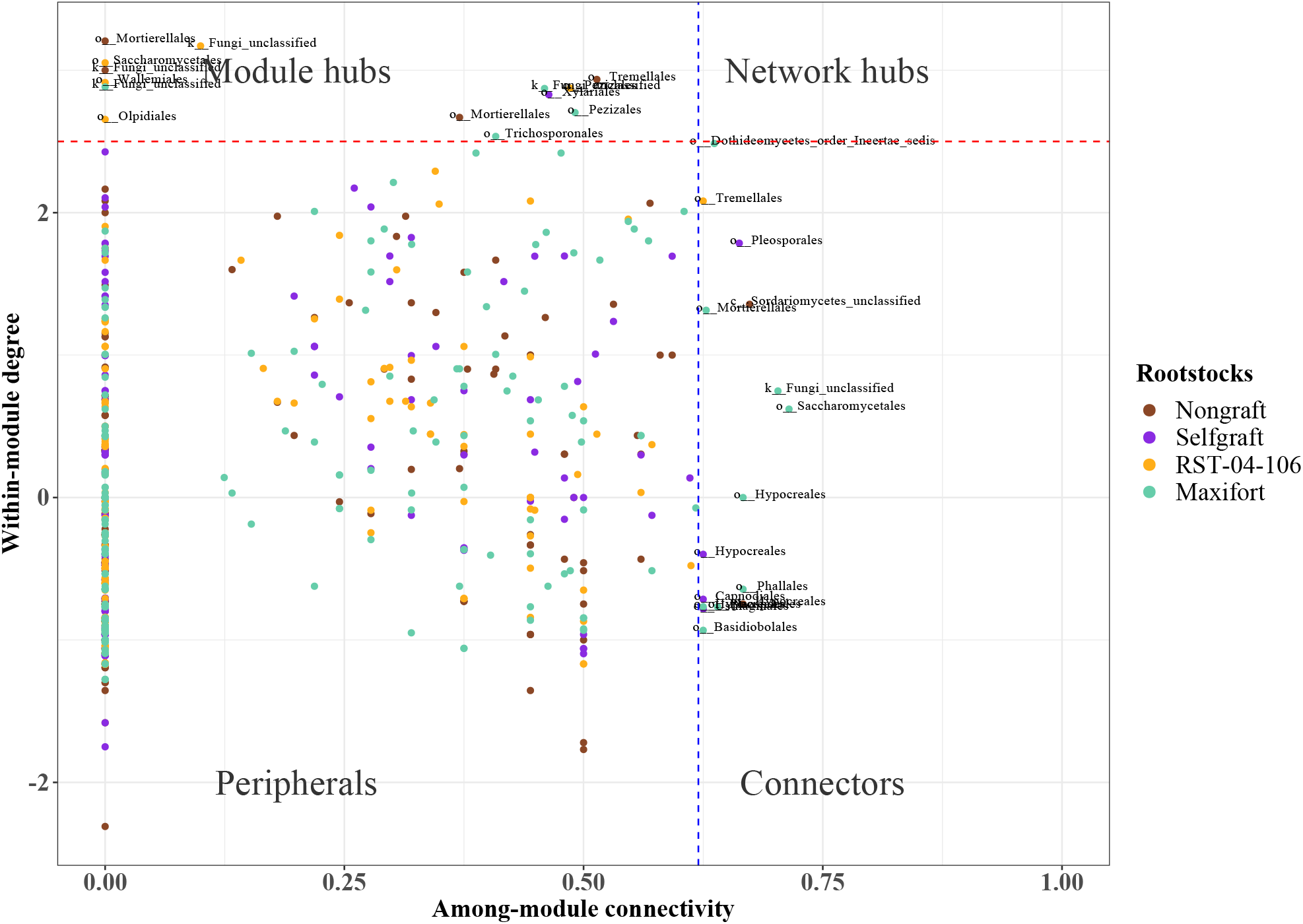
Partitioning of rhizosphere fungal OTUs according to their network roles. Nodes were divided into four categories based on within-module degree and among-module connectivity. The blue dashed line represents a threshold value (0.62) for among-module connectivity, and the red dashed line represents a threshold value (2.5) for within-module degree. Nodes were categorized as peripherals, connectors, module hubs, and network hubs. Node color indicates rootstock treatment (nongraft BHN589, selfgraft BHN589, and BHN589 grafted on two hybrid rootstocks (RST-04-106 and Maxifort)).

### Lasso regression, GLM, and PhONA

Using lasso regression and GLM models, we identified the OTUs predictive of tomato yield in each compartment in each rootstock. The number of predictive OTUs identified by the varImp function was about the same across the rootstock treatments in both the compartments. However, not all the predicted OTUs were associated with other OTUs in the network models. The Maxifort rhizosphere had the highest number of OTUs (10) associated with other OTUs in the network models. Only a subset of the entire microbiome was predictive of the yield, among which only a few microbes were associated with other microbes in the network models.

## DISCUSSION

This study demonstrated the effect of rootstocks on RAF community composition and structure. General diversity-based analyses indicated a rootstock effect. The most productive hybrid rootstock, Maxifort, supported higher fungal richness and Shannon entropy, as well as a greater number of DAOTUs than the controls, consistent with the expectation that higher diversity and a higher number of responsive taxa (DAOTUs) would be associated with a more productive genotype. Also consistent with our expectations, we observed higher microbial diversity and fewer responsive taxa (DAOTUs) in the rhizosphere compared to the endosphere. The integrated host phenotype and OTU network in the PhONA identified potential candidate taxa for each rootstock, and community structures in the endosphere and rhizosphere compartments. The general network analysis found more interactions and more complex network structures in fungal communities associated with Maxifort, consistent with our expectation that a more productive rootstock would have greater community complexity. Community complexity, when defined in terms of mean node degree, differed between the root compartments: the endosphere community was less complex than the rhizosphere community in all the rootstocks. Overall, our study i) showed that rootstocks and grafting are significant drivers of RAF community composition, diversity and structure, and ii) introduced and illustrated the use of PhONA as an analytical framework to select potential candidates for microbiome-based agriculture. Potential candidates are those taxa that were directly predictive of higher yield (taxa with direct positive links with the yield node in the network), taxa that have positive associations with taxa positively associated with yield, and/or taxa that have negative associations with taxa negatively associated with yield. There is the potential to consider taxa multiple steps removed from the yield response, with the understanding that uncertainty about the link to yield increases the more steps the taxon is from the yield node. From a practical standpoint, our results indicate the potential for using plant genotypes and agricultural practices to modulate plant-associated microbial communities, and the potential for the PhONA framework to improve identification of candidate taxa to support microbiome-based crop production.

In contrast to network models that portray only microbe-microbe interactions, PhONA integrates the results of GLM models of microbe association with phenotypic traits to support inference about candidate taxa and predictive microbiome analyses. Thus, candidate taxa can be selected not only because they have a direct association with the host response variable(s), but also because they are indirectly associated with the host response variable through their associations with community members that have direct associations with host traits. For instance, a node that has a positive association with the system phenotype node (in our case, yield) might have negative or positive associations with other OTUs. Such OTUs with indirect positive associations with the desired phenotype might also be included in biofertilizer consortia. Using a PhONA, a rational consortium can be selected based on the phenotype of interest. Applying PhONA for disease or pathogen resistance phenotypes could be useful for designing rational biocontrol consortia. We also observed some OTUs with direct negative associations with the yield node. Efforts to control negatively associated taxa, as well as the taxa that have positive associations with these taxa, might contribute to maximizing yield. Although we did not observe any disease symptoms in our experiments, OTUs negatively associated with the yield node might represent a case of asymptomatic negative microbial effects on yield. Efforts to explore negatively associated OTUs might provide opportunities to minimize asymptomatic yield loss in crops.

The main goal of PhONA is to provide a systems framework to generate hypotheses about the role of microbiome components in host function and performance, and to support the potential for mechanistic/predictive models to better understand host-microbiome interactions. *In planta* experiments with fungal cultures are essential to test the hypotheses generated by these models, to help to differentiate between associations that are based on consistent biological interactions and not simply based on shared (or opposing) environmental niche preferences. It is important to be cautious in attributing biological interactions to the key structures in network attributes because the links in the PhONA may or may not depict biological interactions. That is, many links may represent only correlative relationships and not causal ones (28). For instance, the hub node in the network is often regarded as a key node, but the high number of links with the hub node in the association network could be due to shared niches, biological interactions, or a mixture thereof. If the associations are mostly due to shared niches, removing such a hub node will have a more limited effect, whereas removal of a hub node involved in many biological interactions could lead to significant effects on the microbial community.

RAF community composition, diversity, and interactions differed between the endosphere and rhizosphere compartments. Although the endosphere and the rhizosphere are physically adjacent, they are distinct in community composition and diversity. Compartment specificity in community composition and diversity has been reported for other plant species, in both natural and agricultural settings (31, 69, 70). Usually, bulk soil is considered a source of microbial communities, a subset of which is selected for in the rhizosphere (31, 71), mainly as a function of root exudates and rhizodeposits (43, 72–75). Selection of the rhizosphere microbiome could be specific (*e.g*., antagonistic to plant pathogens) (76, 77) or more general with less influence of host genotype. In comparison, the endosphere of host plants often supports lower microbial diversity compared to the rhizosphere (70, 71). Host tissues and defense systems act as biotic filters (2). As a result, the microbiome is more specialized in the endosphere than in the rhizosphere. RAF compartment specificity may also be an important consideration for microbe-based disease management strategies – especially for the management of pathogens or pests that are compartment specific, such as endoparasites and ectoparasites.

RAF community composition and diversity also differed among the rootstock genotypes. Plant genotypes can structure root-associated fungal communities (31, 70, 78). The commercial rootstocks in our study have been bred to provide resistance against specific soilborne pathogens and pests. Small host genotypic differences could alter the physiological and immunological responses in the root systems, thereby selecting genotype-specific RAF communities (79). For example, some root exudates and metabolites could be specific to a plant genotype (80–82) and provide specific control of microbial communities (83–85). In some cases, the host genotype effect can be directly attributed to root anatomy (77, 85, 86). Efficient root types and architectures are desired agronomic traits to cope with biotic and abiotic stresses (87), and root systems vary among and within plant species (86). Moreover, the effect of plant genotypes on microbial communities in the root system may be linked to the flow of nutrients between the aboveground-scion and belowground-rootstocks, where vigorous rootstock genotypes could drive greater resources to the microbial communities by supporting larger scion biomass. In such a positive nutrient feedback between the scion and rootstock, rootstock genotype appears to be a more critical driver than the scion genotype (88). Rootstock genotypes supporting higher yield and biomass may support higher microbial diversity by excreting a greater volume of photosynthates as root exudates and metabolites. Although we did not evaluate root exudates, and used the same scion across the study, our study is consistent with a role of higher yield and biomass (as for the Maxifort rootstock) being associated with higher fungal diversity. Additionally, we observed an effect of rootstock on the RAF community composition. Collectively, the results support our expectations of rootstock-specific control of the RAF community.

Our definition of complexity is based on interactions in networks, using a definition similar to that used in other microbiome network analyses (25, 34, 89). However, a greater number of interactions and complex network structures/motifs would tend to be observed whenever more nodes exist in these association networks, an inherent relationship not always considered in studies of complexity in microbiomes. The higher number of OTUs associated with Maxifort would tend to result in higher complexity compared to rootstocks with fewer OTUs. Another potential measure of complexity is network density, the proportion of links observed in a network relative to the total number of possible links. For all the rootstocks we studied, network density was similar (0.04) in both compartments, indicating similar community complexity. Statistical methods comparable to rarefaction, designed to equalize the number of nodes across networks or methods to balance OTU richness for sampling efforts (90), will be a valuable future effort for understanding how network complexity responds to treatments and for making comparisons across studies. In addition, methods to optimize and automate the selection of association thresholds to define the pairwise relationships in a microbiome network is a gap and opportunity for improving microbiome network analyses. For graphics in the figures in this analysis, we selected a level of association such that an interpretable number of OTUs were depicted for visual consideration. Studies directly applied to identify potential microbial assemblages for agricultural applications could benefit from exploring results for a range of thresholds.

Our study indicates the rootstock-genotype specific effect on RAF diversity, composition, and interactions, and also demonstrates integration of system phenotypes such as plant yield in a network-based model to support selection of candidate taxa for biological use. However, in sequence-based studies such as ours, the biological and functional significance of the candidate OTUs remains unknown. Follow-up experiments with fungal cultures will be necessary to determine the biological roles of the candidate OTUs, and to differentiate causal associations from correlations based on niche preference. Similarly, further development of PhONA to incorporate temporal microbiome data and Bayesian learning and inference methodologies (91, 92) has the potential to support causal inference, including understanding of directionality in microbiome networks. PhONA utilizes a lasso regression and GLM to link OTUs with a system phenotype, although many other models such as random forest and other machine learning approaches (93) could also be employed. Given the nature of microbiome data, having a high number of features (p) and relatively small number of samples (n), other models to address the n x p problem can improve PhONA predictions. Rather than pure prediction, our methods aim to find the key predictors and use them in the GLM model for evaluating associations with the yield response. PhONA focused on finding the attributable predictors/OTUs that are key to biological interventions, which are missed in approaches that are focused purely on prediction (94). Smaller sample size was a limitation in our current study, reflecting the challenge of processing a large number of plant replicates, and we did not validate the results from our model by splitting data into training and test sets. A rigorous model validation step would improve the accuracy of PhONA. As lab-based technologies and computational resources become less expensive, studies with large sample sizes are becoming more practical and, when combined with an analytical framework like PhONA, microbial community analyses can go beyond simple analyses of diversity to help make microbiome-based agriculture a reality.

## Acknowledgements

We appreciate support from the Ceres Trust, USDA NCR SARE Research and Education Grant LNC13-355, Foundation for Food and Agriculture Research Grant FF-NIA19-0000000050, the Kansas Agricultural Experiment Station, the National Institute for Mathematical and Biological Synthesis (NIMBioS), and the University of Florida. We thank members of the Garrett lab for their comments on an early version of the manuscript. R.P., K.A.G., A.J., M.M.K., and C.L.R. designed the study. R.P. and L.G.M. collected and processed the samples. R.P. analyzed and interpreted the data. R.P., K.A.G., and A.J. wrote the manuscript. R.P., K.A.G., A.J., M.M.K., C.L.R., and L.G.M. revised the manuscript. All authors read and approved the final manuscript.

---

**FIG S1.**
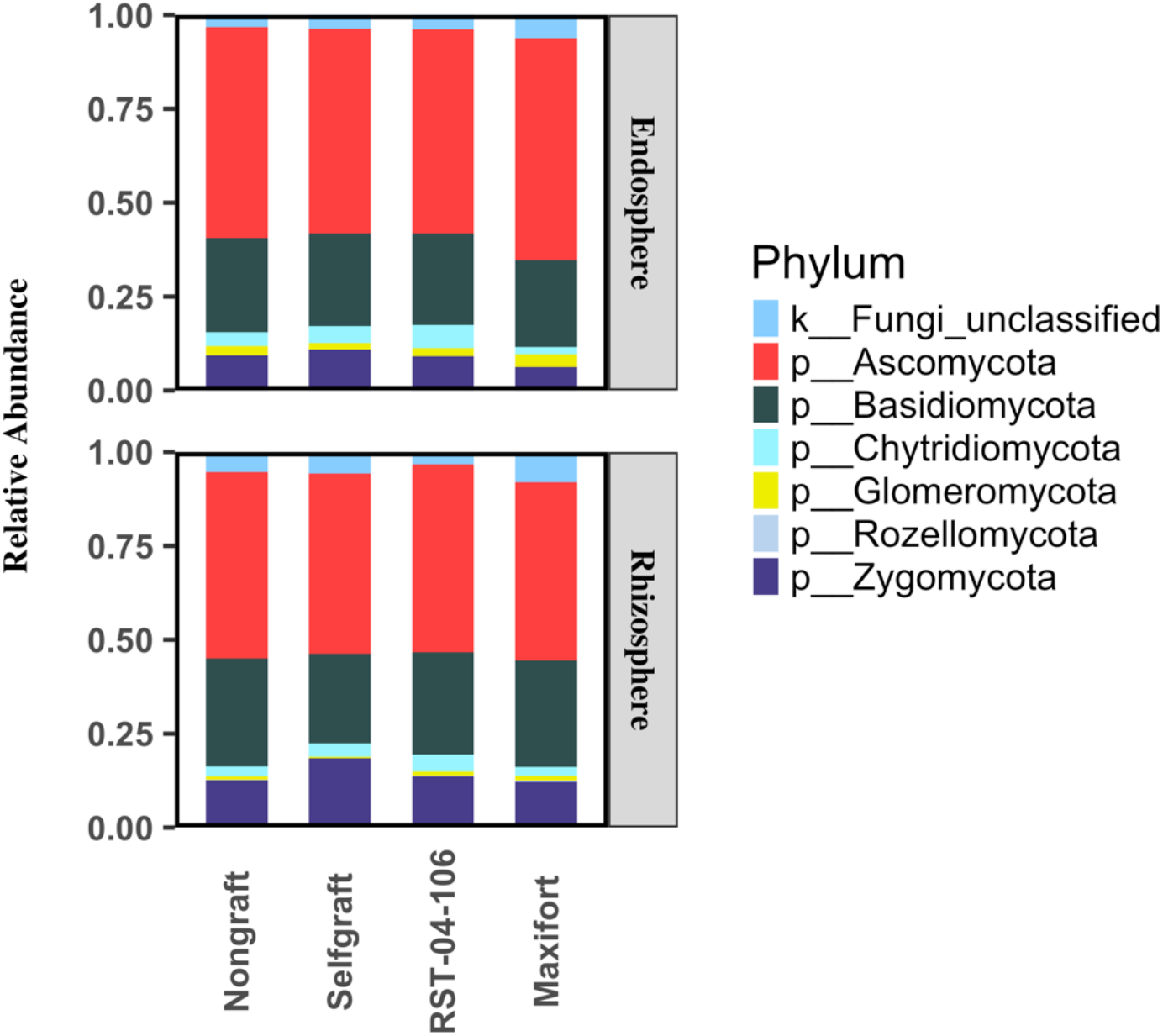
Relative abundance of endosphere and rhizosphere fungi at the phylum level recovered from four tomato rootstock treatments: nongraft BHN589, selfgraft BHN589, and BHN589 grafted on two hybrid rootstocks (RST-04-106 and Maxifort). Each individual bar represents a rootstock treatment, and the colored area within the bar represents the relative abundance of the corresponding phylum.

**FIG S2.**
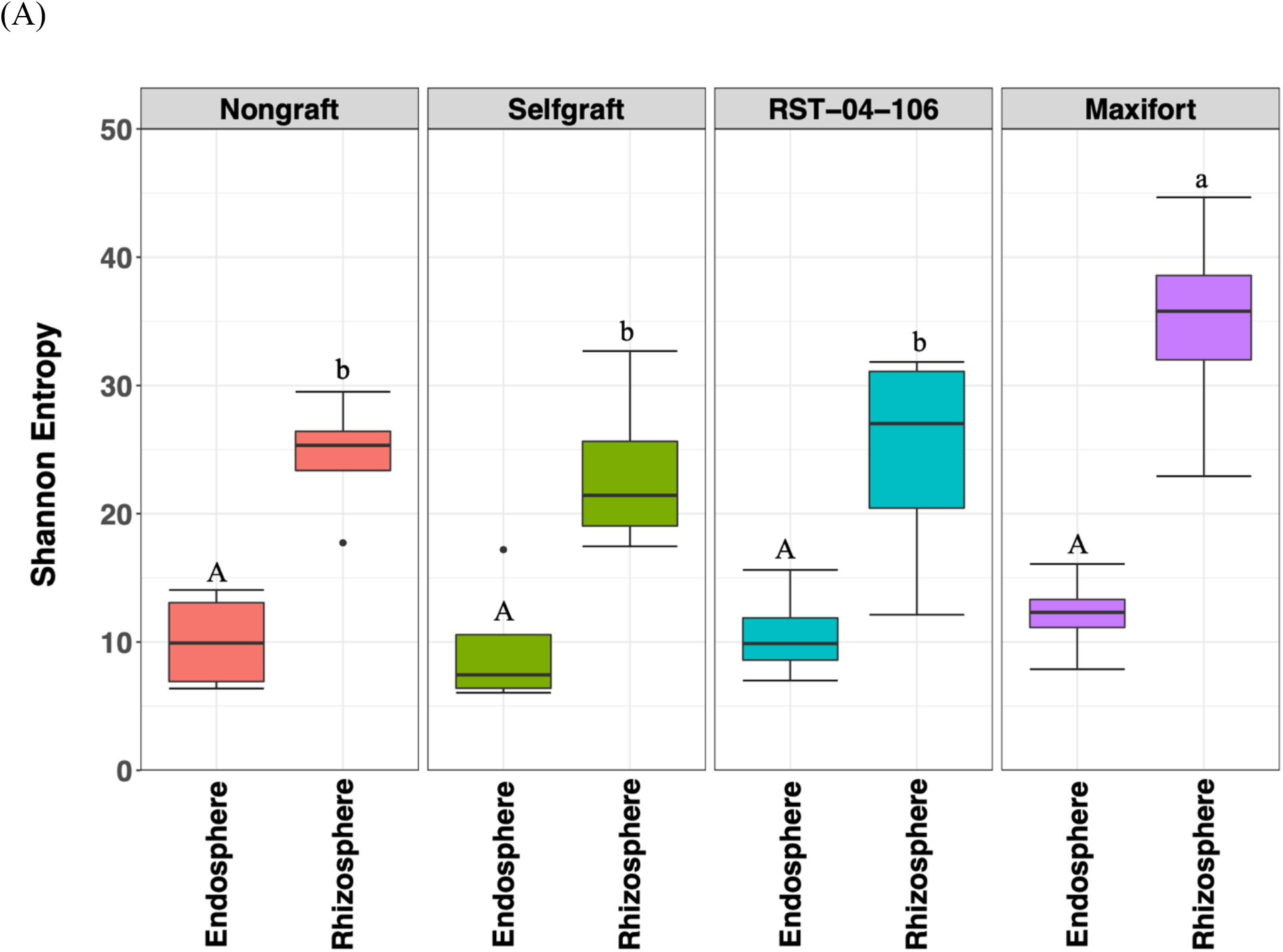
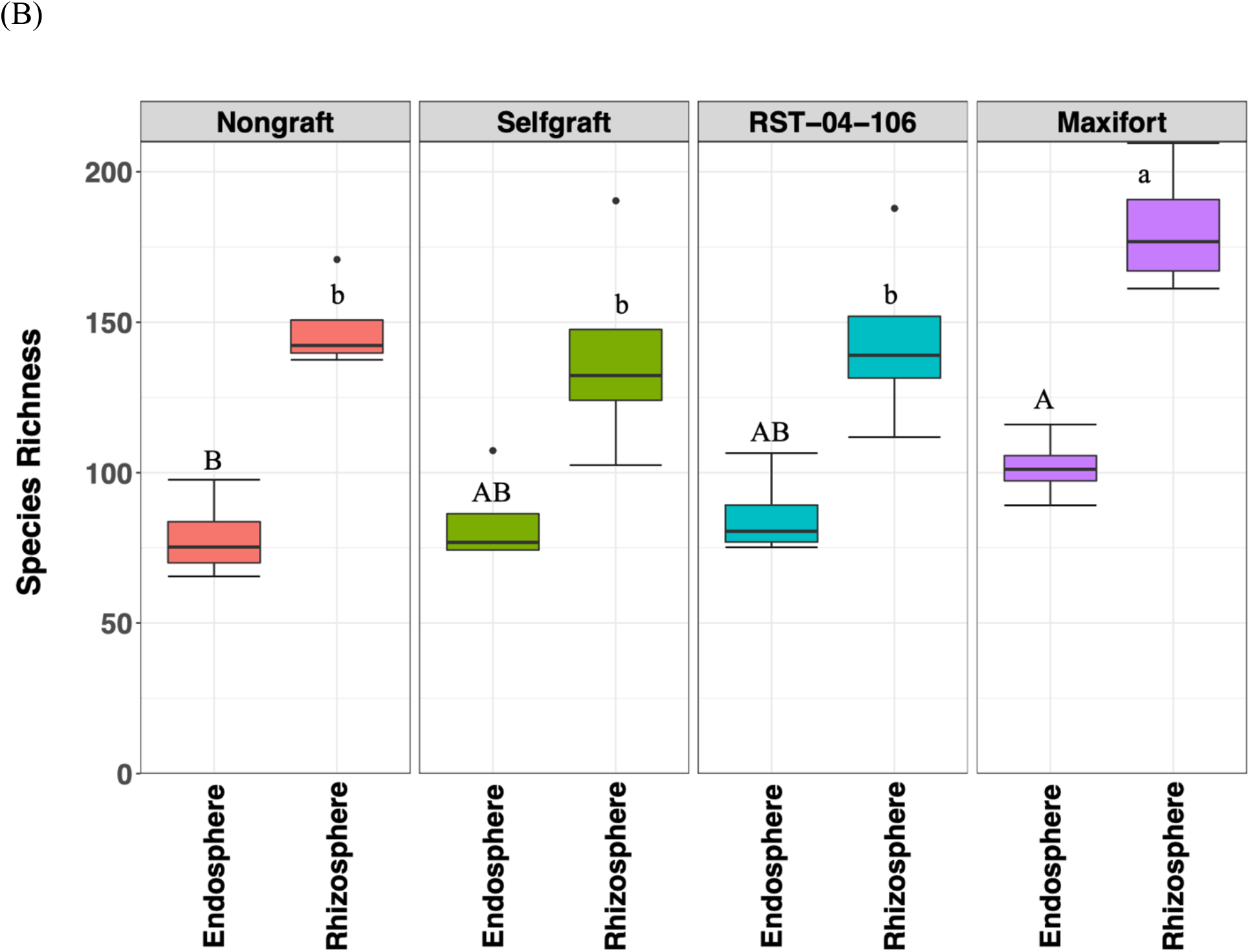
Comparison of overall fungal diversity (A) and richness (B) associated with tomato rootstock genotypes and controls, evaluated in the endosphere and rhizosphere. The plot is divided by the four tomato rootstock treatments: nongraft BHN589, selfgraft BHN589, and BHN589 grafted on two hybrid rootstocks (RST-04-106 and Maxifort). Shannon entropy and species richness, measures of community diversity, were both higher for Maxifort (p < 0.005) compared to the self-graft and RST-04-106 in the rhizosphere, while there was not evidence for a difference in Shannon entropy in the endosphere (p = 0.634). Treatment means were separated using the “difflsmeans” function as specified in the lmerTest package in R. Tests for boxplots sharing a letter or letter case type had p > 0.05.

**FIG S3.**
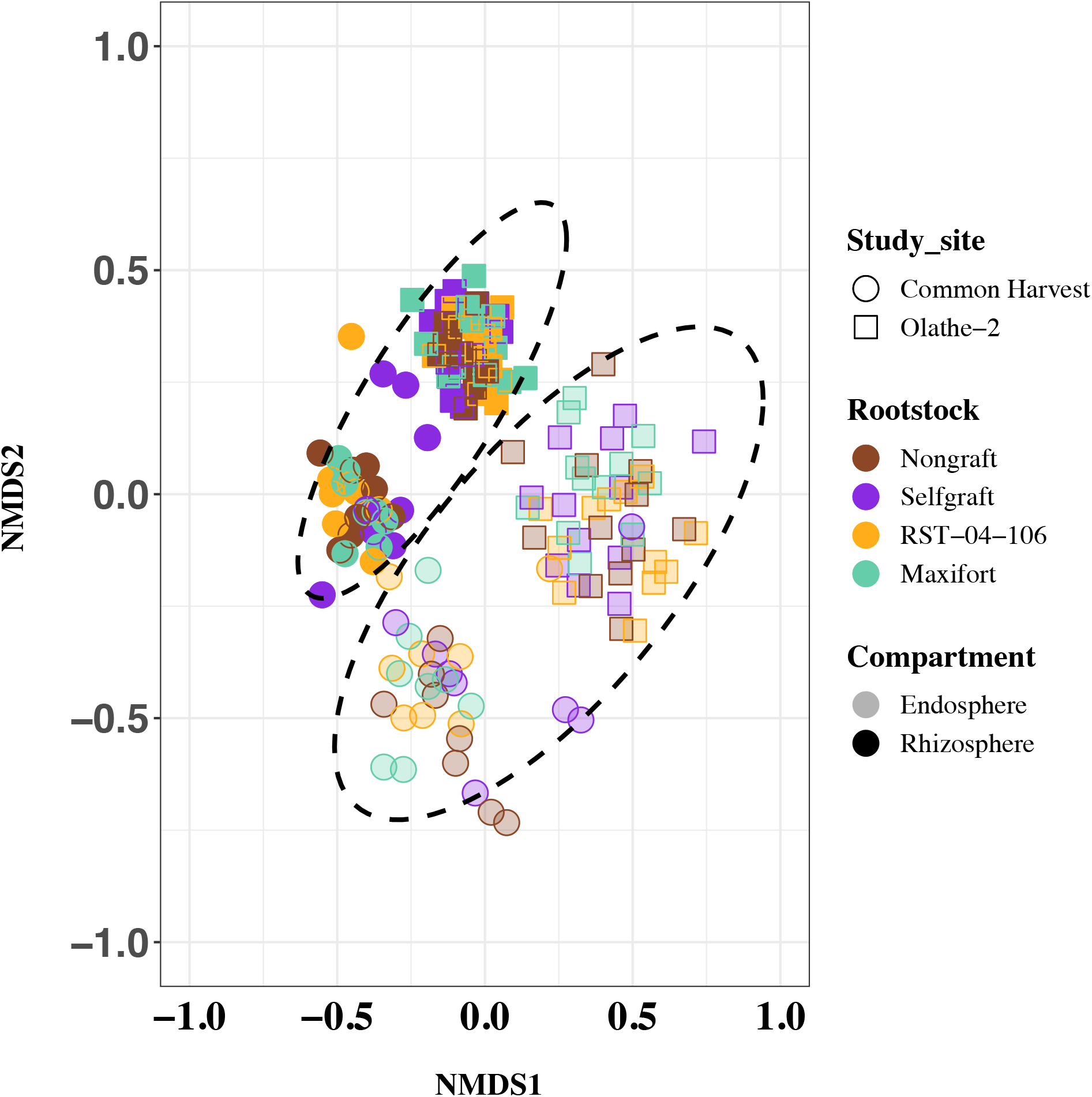
Non-metric multidimensional scaling (NMDS) ordination plot of samples labeled by tomato rootstock (nongraft BHN589, selfgraft BHN589, and BHN589 grafted on two hybrid rootstocks (RST-04-106 and Maxifort)), compartment (endosphere or rhizosphere), and study site, based on the Bray-Curtis dissimilarity distance matrix of fungal OTUs. Color indicates rootstock treatment, shape indicates study site, and size indicates compartment. Ellipses surrounding the samples indicate the 95% CI of the endosphere and rhizosphere sample centroids.

**FIG S4.**
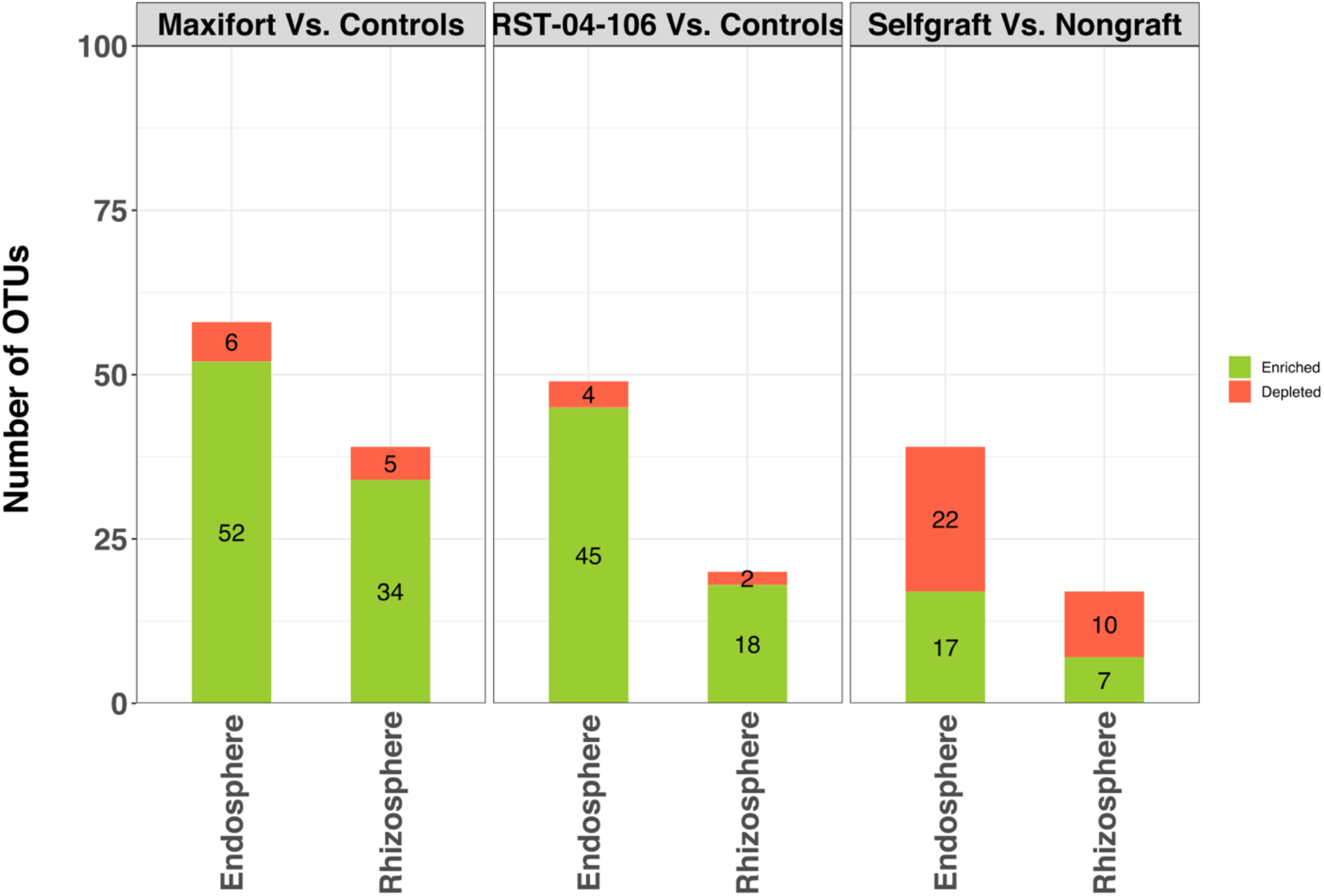
Number of DAOTUs in a contrast analysis, evaluated for the endosphere and rhizosphere compartments for four tomato rootstock treatments: nongraft and selfgraft BHN589, and BHN589 grafted on two hybrid rootstocks (Maxifort and RST-04-106). The green color in each bar represents the number of enriched taxa, and the red color represents the number of depleted taxa. The number of differentially changed taxa was greater for the endosphere than for the rhizosphere. Among the contrast pairs, hybrid rootstocks had a greater number of enriched taxa compared to depleted taxa. However, the number of depleted taxa was higher compared to enriched taxa in the controls. Among the treatments, Maxifort had the highest number of DAOTUs in both compartments.

**FIG S5.**
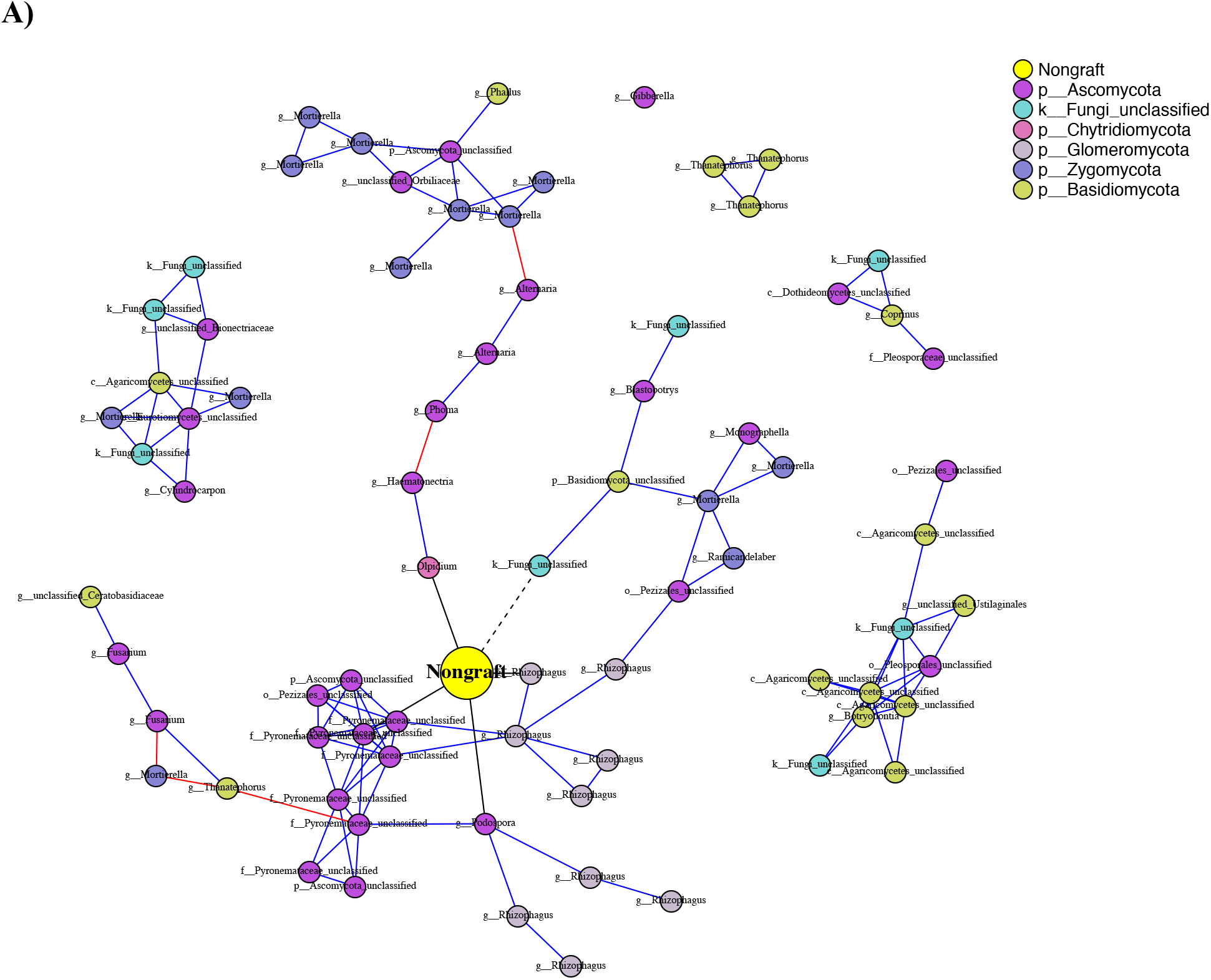
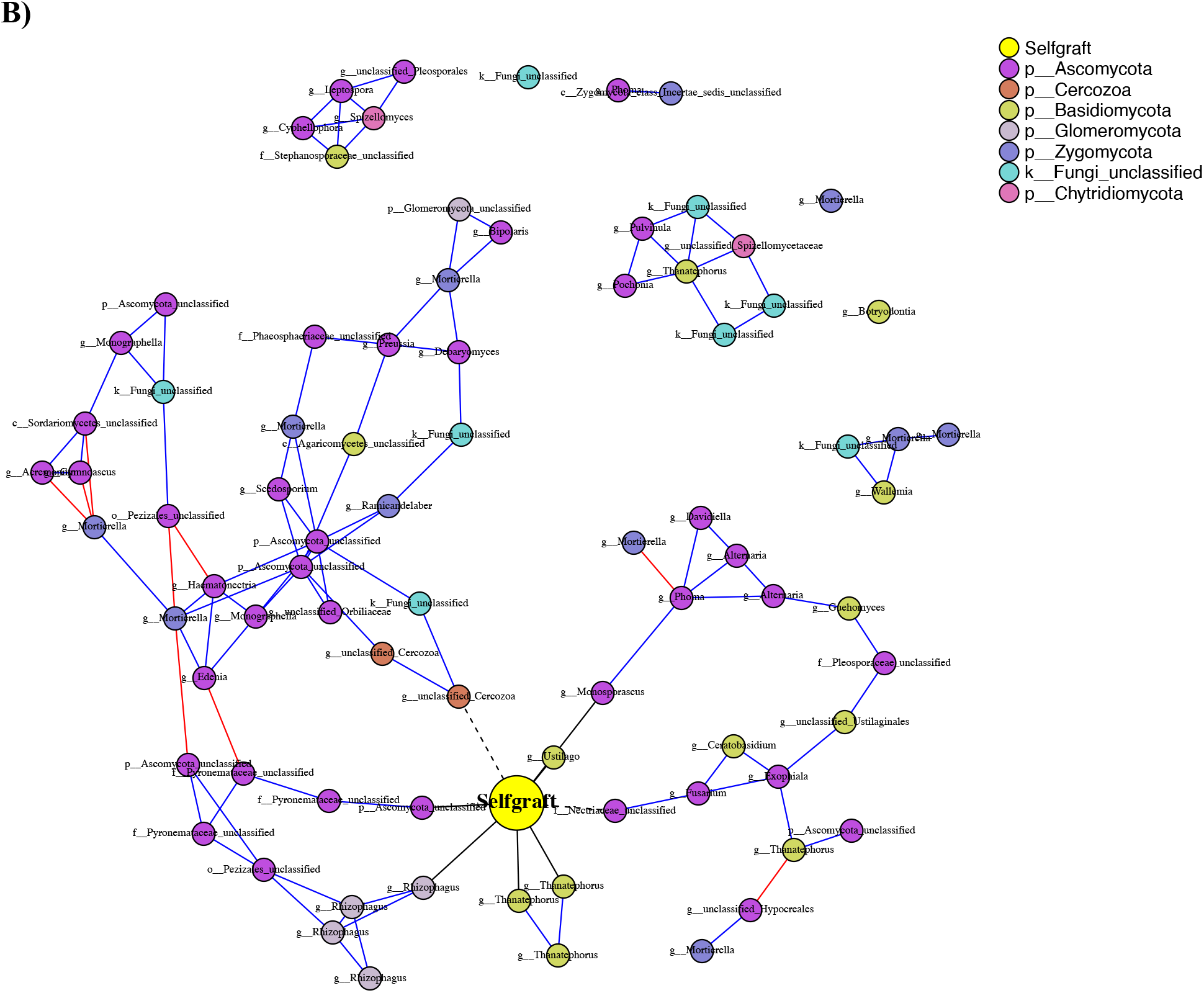
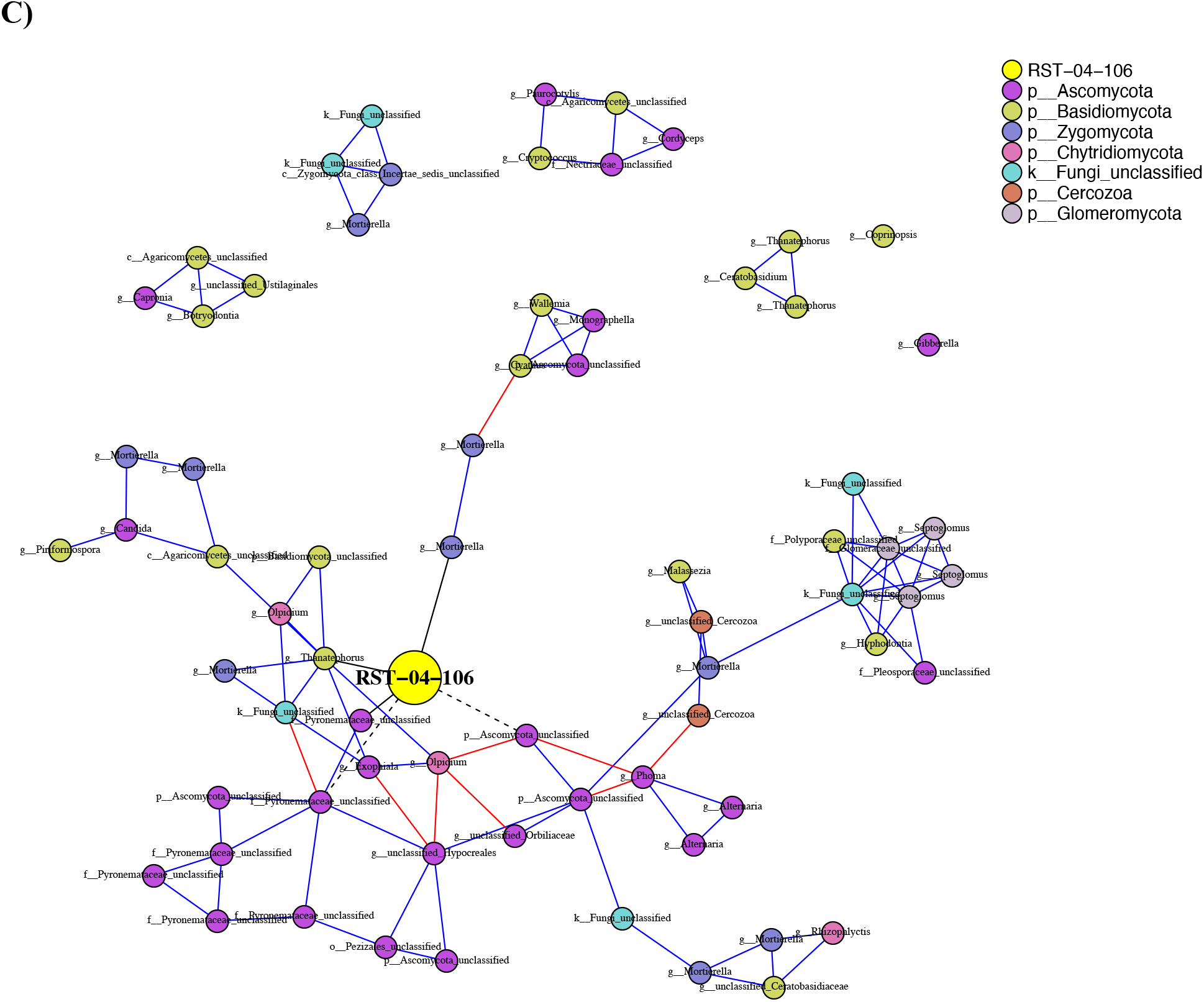
Phenotype-OTU network analysis (PhONA) of endosphere fungal taxa for tomato rootstock treatments: (A) nongraft and (B) selfgraft BHN589, and (C) BHN589 grafted on RST-04-106. Node color indicates the phylum, except that the yellow-color node represents yield associated with the rootstock. Nodes connected to the rootstock yield node with black links are taxa that were predictive of rootstock yield, where dotted and solid lines indicate negative and positive associations with the yield node, respectively. Red and blue links represent negative and positive associations, respectively, between OTUs. Nodes are labeled with the finest-resolution taxonomic categorization available.

**FIG S6.**
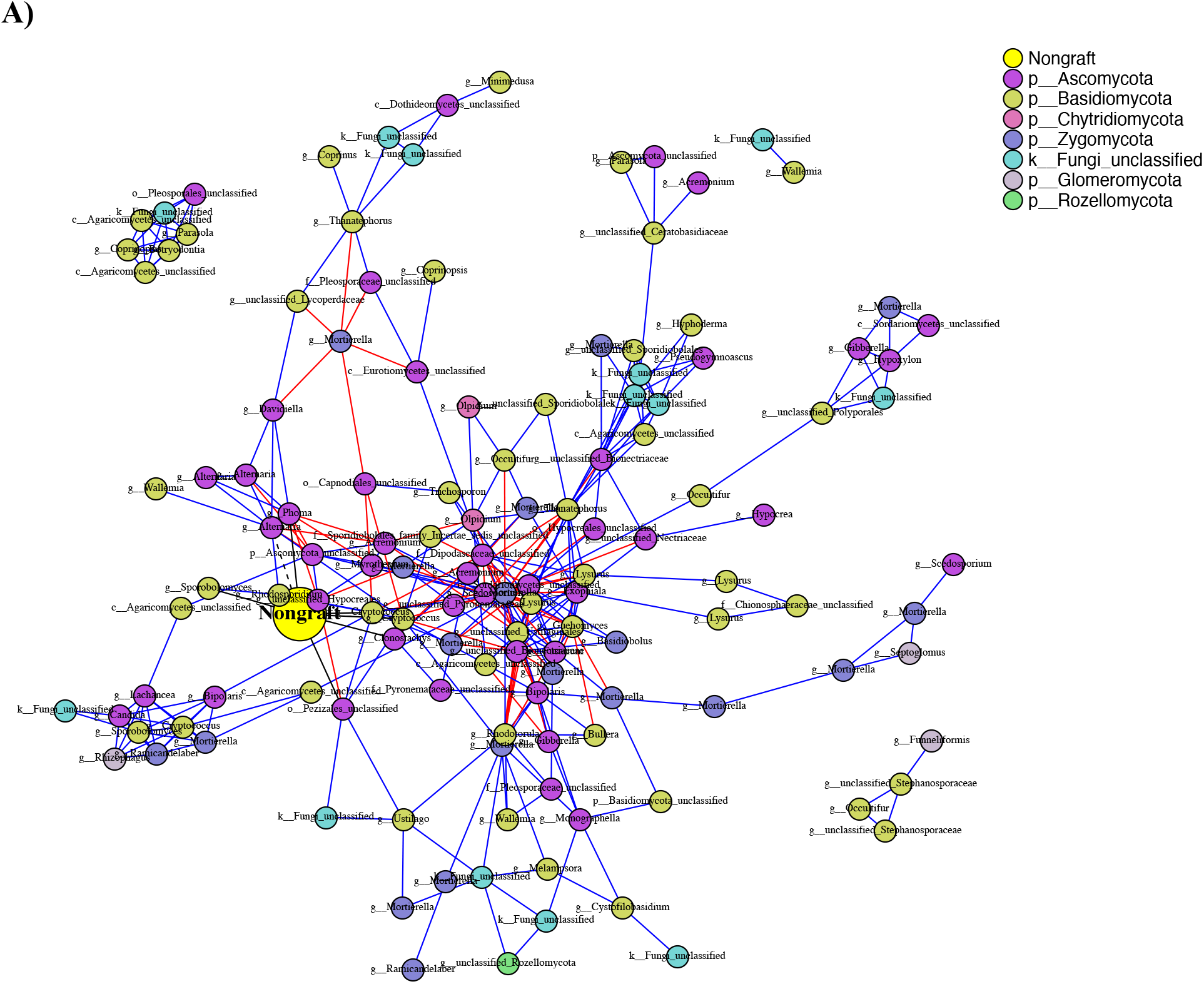
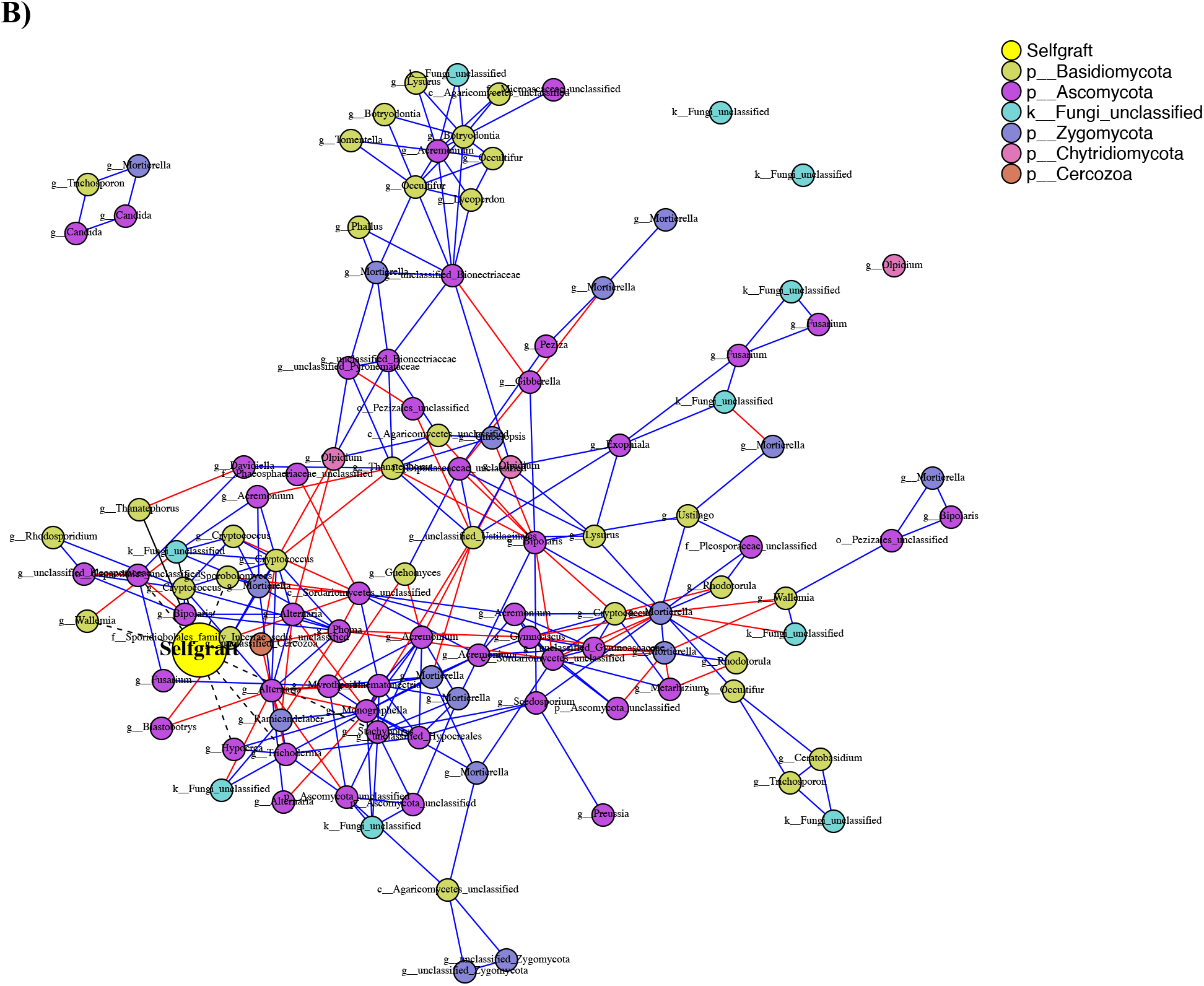
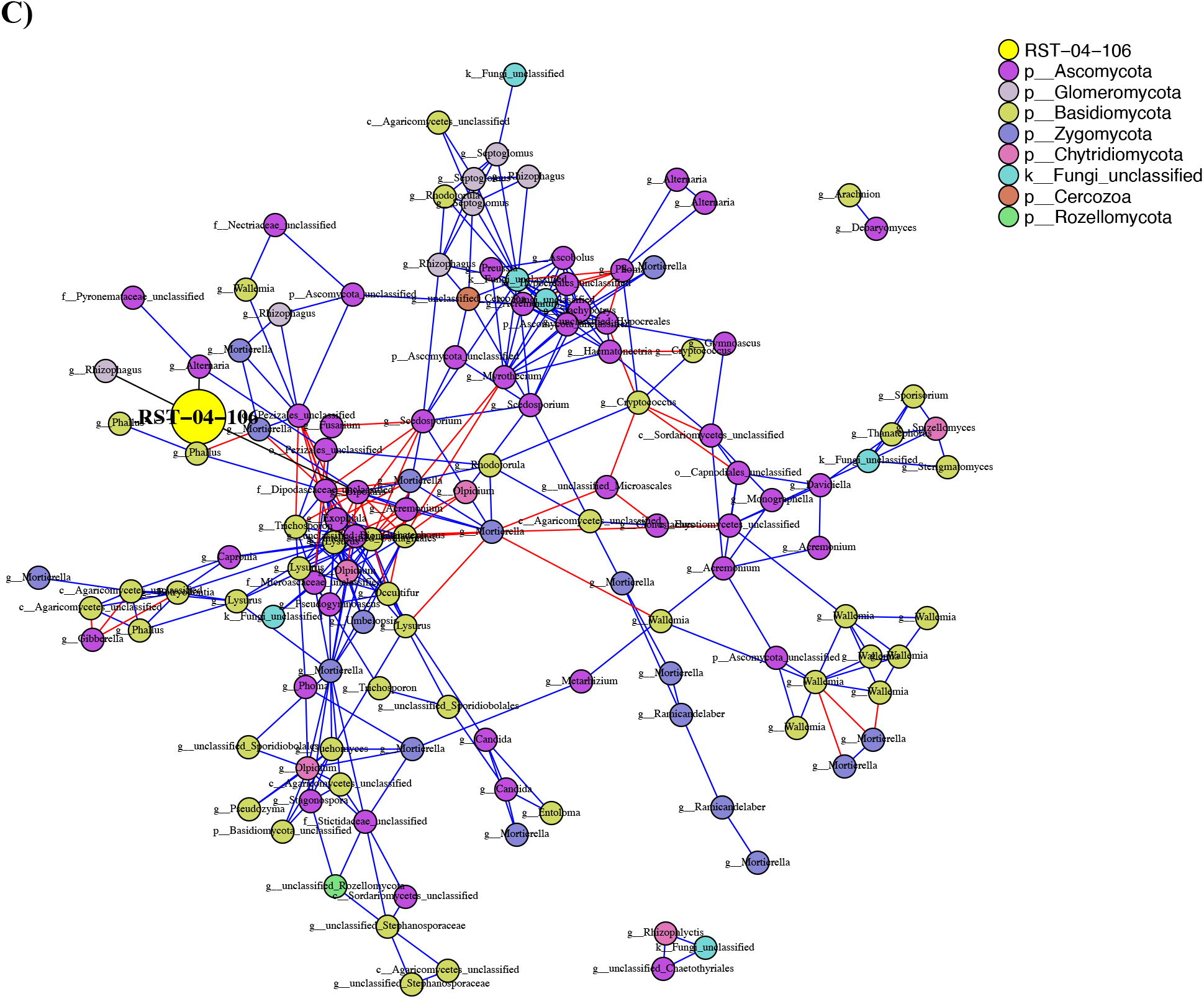
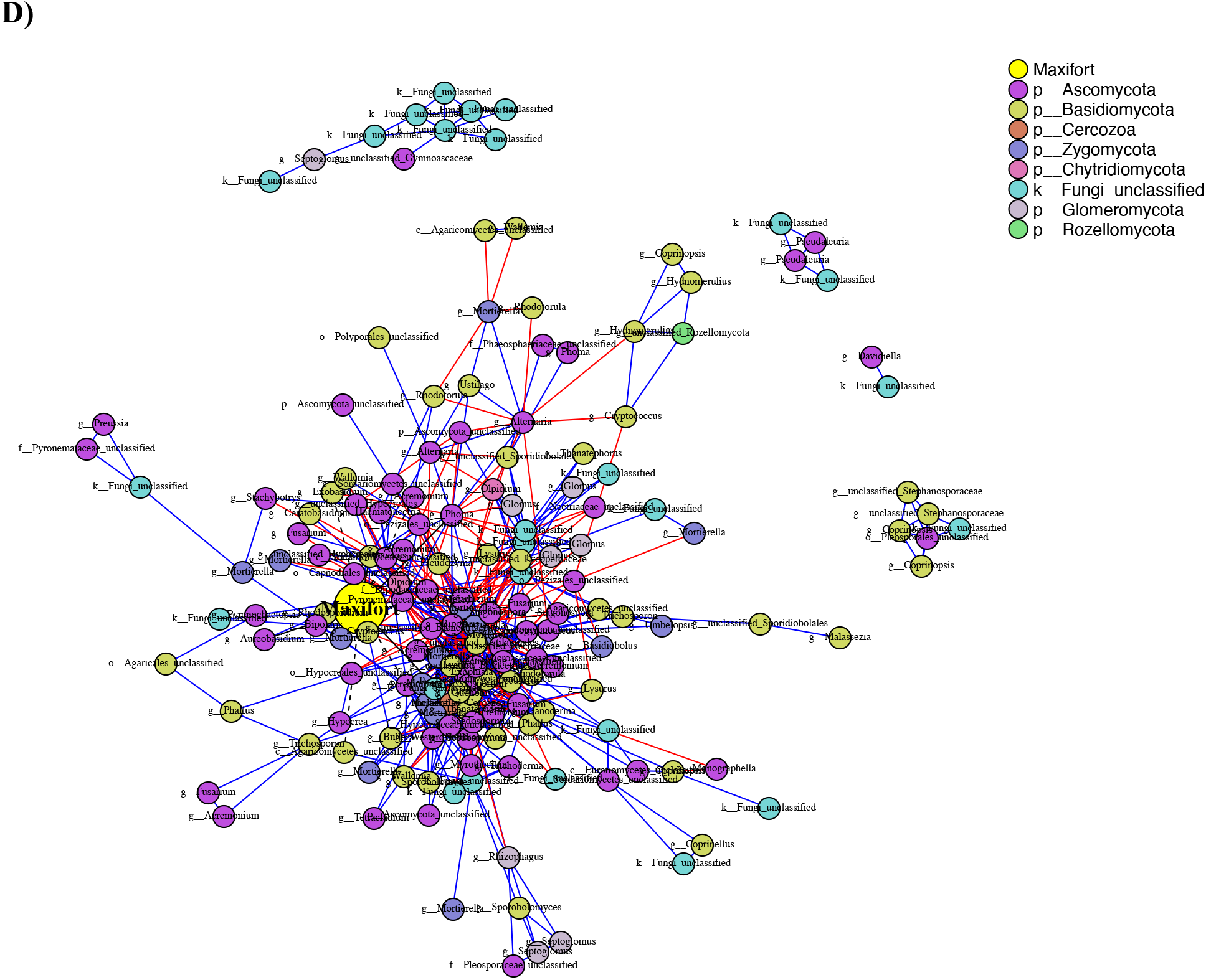
Phenotype-OTU network analysis (PhONA) of rhizosphere fungal taxa for tomato rootstock treatments: (A) nongraft and (B) selfgraft BHN589, and BHN589 grafted on two hybrid rootstocks ((C) RST-04-106 and (D) Maxifort). Node color indicates the phylum, except that the tomato-color node represents yield associated with the rootstock. Nodes connected to the rootstock yield node with black links are taxa that were predictive of rootstock yield, where dotted and solid lines indicate negative and positive associations with the yield node, respectively. Red and blue links represent negative and positive associations, respectively, between OTUs. Nodes are labeled with the finest-resolution taxonomic categorization available.

**TABLE S1.** Sites included in the study, their soil type, and geographic coordinates.

| Study sites | Location | Latitude | Longitude | Soil type |
| --- | --- | --- | --- | --- |
| Olathe Horticulture Research and Extension Center (OHREC) | Johnson County, KS | 38.88N | 94.99W | Chase silt loam |
| Common Harvest | Douglas County, KS | 38.96N | 95.20W | Eudora-Kimo complex |

**TABLE S2.** Results of the multivariate permutational analysis of variance (PERMANOVA) for fungal taxon abundance data. Permutation was based on the Bray-Curtis distance matrix generated for root associated fungal communities at the OTU level from four tomato rootstock treatments: nongraft and selfgraft BHN589, and BHN589 grafted on two hybrid rootstocks (Maxifort and RST-04-106) (1000 permutations). P values < 0.05 are in bold.

| Factor | Sum of Squares | % Explained | P value |
| --- | --- | --- | --- |
| Rootstocks | 1.24 | 2.07 | < **0.01** |
| Compartment | 5.36 | 8.92 | < **0.001** |
| Study_Site | 5.01 | 8.34 | < **0.001** |
| Year | 3.23 | 5.38 | < **0.001** |
| Rootstocks:Compartment | 0.58 | 0.97 | 0.972 |
| Rootstocks:Study_Site | 1.39 | 2.32 | < **0.01** |
| Compartment:Study_Site | 1.59 | 2.65 | < **0.001** |
| Rootstocks:Year | 1.00 | 1.66 | 0.111 |
| Compartment:Year | 1.00 | 1.66 | < **0.001** |
| Study_Site:Year | 1.46 | 2.43 | < **0.001** |
| Rootstocks:Compartment:Study_Site | 0.55 | 0.91 | 0.998 |
| Rootstocks:Compartment:Year | 0.49 | 0.82 | 1.000 |
| Rootstocks:Study_Site:Year | 1.22 | 2.03 | < **0.01** |
| Compartment:Study_Site:Year | 0.60 | 1.00 | < **0.01** |
| Rootstocks:Compartment:Study_Site:Year | 0.60 | 1.00 | 0.989 |
| Residuals | 34.75 | 57.86 |  |
| Total | 60.07 | 100.01 |  |

**TABLE S3.**
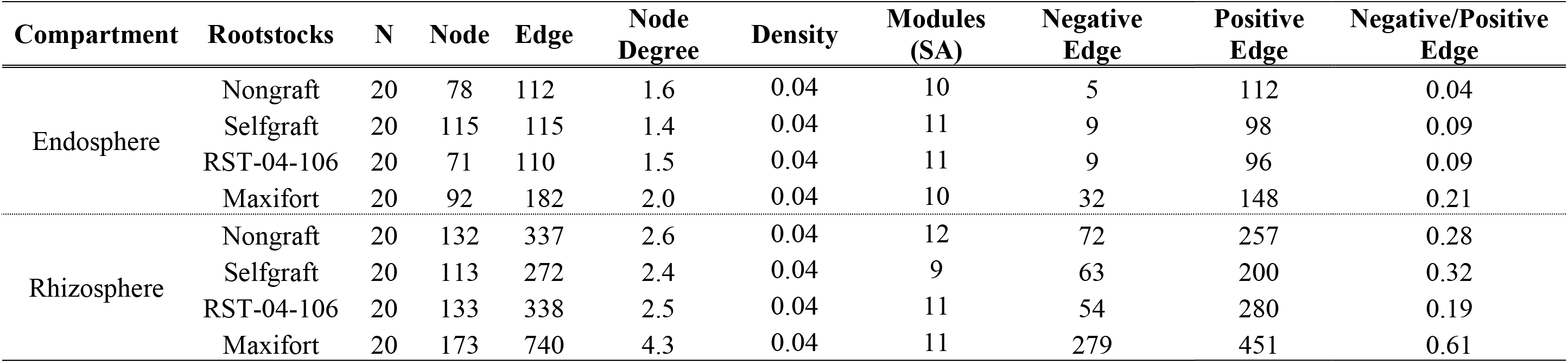
Network attributes and links observed in the fungal association networks for four tomato rootstock treatments: nongraft and selfgraft BHN589, and BHN589 grafted on two hybrid rootstocks (Maxifort and RST-04-106) in each of rhizosphere and endosphere compartments.

